# Neuron-intrinsic and glial pathways regulate sensory cilia regeneration in adult *C. elegans*

**DOI:** 10.64898/2026.05.15.725580

**Authors:** Kirsten Judge, Alison Philbrook, Stephen Nurrish, Samantha Leslie, Laura Grunenkovaite, Yu-Ming Lu, Piali Sengupta

## Abstract

Primary cilia are microtubule-based organelles that mediate cellular responses to environmental cues. Although cilia disassemble and reassemble upon cell cycle entry and exit, respectively, in dividing cells, it remains unclear whether postmitotic cells such as neurons can regenerate these structures *in vivo* following injury to restore function, and whether this process recapitulates developmental ciliogenesis. Here we show that a subset of sensory neuron cilia in adult *C. elegans* regrows following conditional truncation and restores neuronal functions. This regeneration is regulated by both cell-intrinsic and extrinsic mechanisms that are in part distinct from those employed during embryonic ciliogenesis. We find that the conserved ciliogenic DAF-19 RFX transcription factor is dispensable for cilia maintenance in the adult but is required for regeneration, in part via transcriptional upregulation of a subset of ciliary intraflagellar transport (IFT) genes. We further identify the DLK-1 dual leucine-zipper kinase and the CEBP-1 C/EBP transcription factor, previously implicated in axon regeneration, as necessary for efficient cilia regrowth but not for developmental ciliogenesis. We show that *cebp-1* expression is induced during cilia truncation and regrowth and that this induction is also DAF-19-dependent. Finally, we show that cilia truncation and regeneration dynamics vary in a neuron type-specific manner and are modulated by signals from surrounding glia. Our results establish that the cilia of mature neurons can regenerate and recover functions *in vivo* and identify conserved pathways that regulate this process in adult animals.

## INTRODUCTION

Primary cilia are microtubule-based organelles that project from the membranes of most cell types and transduce environmental signals to regulate cellular and organismal functions ^1,2^. Consequently, loss of cilia or disruption of ciliary signaling underlies a broad class of disorders termed ciliopathies ^3,4^. Given the physical location of cilia on the cell surface, these organelles are particularly vulnerable to damage by chemical or physical agents ^5–8^. Cilia on airway epithelial cells can be injured by airborne pollutants, cilia on endothelial cells are disassembled in response to high laminar shear stress, and cilia on diverse cell types can be damaged upon viral infection ^5,7,9–12^. In dividing cells, damaged cilia can be replaced by highly regulated cycles of ciliary disassembly and reassembly correlated with cell-cycle entry and exit, respectively ^13,14^. However, whether cilia loss results in irreversible cellular dysfunction in postmitotic cells such as neurons, or whether cilia of different neuron types can regenerate and restore cellular function in the adult *in vivo* remains to be fully understood.

Cilia regeneration following injury or truncation has been described in multiciliated tissues, cultured cells, olfactory epithelia, the mammalian inner ear, and in unicellular organisms such as *Chlamydomonas* and *Tetrahymena* ^15–23^. In the mammalian brain, neuronal ciliogenesis occurs during embryonic and early postnatal development and cilia are then maintained through adulthood ^2,24–27^. Thus, if cilia were to regenerate following loss at later postnatal stages, regrowth would occur in neurons with developmental states, and in cellular environments, that differ significantly from those during embryonic development. These distinct internal and external conditions may preclude cilia regeneration in mature neurons or require mechanisms that differ from those driving developmental ciliogenesis. Analogous studies on axon regeneration across species have shown that axon regrowth in the adult relies on mechanisms that are partly distinct from those used for axon outgrowth and guidance during development ^28–31^. Moreover, while peripheral axons in mammals regenerate robustly, central axons exhibit limited regeneration due to both intrinsic and extrinsic factors ^32–35^.

In *C. elegans*, the majority of ciliated sensory neurons including the twelve neuron pairs of the bilateral head amphid organs are born in the embryo ^36^. Ciliogenesis in amphid neurons occurs during embryogenesis and these cilia are maintained throughout the life of these animals ^37–39^. Previous studies have shown that restoring ciliary gene function in the adult in ciliogenesis mutants can rescue cilia morphology, and axonemes truncated by treatment with noxious chemicals regrow ^40^, indicating that a subset of sensory cilia retains the capacity for regrowth in the adult ^39,41,42^. However, the intrinsic and extrinsic mechanisms that promote or inhibit cilia regeneration, and whether morphologically diverse cilia of different neuron types can regenerate in the adult, are unknown.

Here we show that a subset of amphid sensory neuron cilia formed during embryonic development can regenerate and restore neuronal functions in adult *C. elegans* following conditional truncation. We find that the essential and evolutionarily conserved ciliogenic DAF-19 RFX transcription factor ^43^ is dispensable for cilia maintenance in the adult but is required for regeneration in part via upregulating intraflagellar transport (IFT) genes necessary for cilia assembly ^44,45^. We further identify roles for the DLK-1 dual leucine-zipper kinase and CEBP-1 C/EBP transcription factor, previously implicated in axon regeneration ^46,47^, in promoting cilia regrowth but not embryonic ciliogenesis. Similar to IFT genes, *cebp-1* expression is upregulated during cilia truncation and regrowth, and we show that this induction is also mediated by DAF-19. Finally, we find that cilia exhibit neuron-specific temporal dynamics of shortening and regrowth, and that glial signals regulate these processes differentially across sensory neuron types. Together, our results describe a set of cell-intrinsic and extrinsic mechanisms that regulate cilia regeneration in mature sensory neurons, and suggest that manipulation of these pathways may restore neuronal functions following cilia injury or loss in adult animals.

## RESULTS

### Sensory cilia regenerate and restore neuronal function following conditional truncation in adult *C. elegans*

Pioneering studies on axon regeneration in adult *C. elegans* used laser microsurgery to sever individual axons and follow axon regrowth ^48,49^. The cilia of eight sensory neurons in the bilateral head amphid sense organs of *C. elegans* are ∼5-7 μm long and are tightly bundled within a channel formed by glial processes (“channel” cilia) ^50,51^, making it impractical to perform high-throughput laser severing of individual cilia. *unc-70* β-spectrin gene mutants in which axons break spontaneously due to defects in the underlying neuronal membrane cytoskeleton have been used to increase throughput of genetic studies on axon regeneration ^52,53^. We, therefore, sought a genetic strategy to conditionally truncate cilia in adult *C. elegans* to initiate studies on cilia regeneration.

Cilia are built by the conserved process of IFT mediated by anterograde kinesin and retrograde dynein motors that traffic ciliary building blocks and signaling proteins into and out of these organelles ^44,45^. The middle segments of channel cilia in *C. elegans* are built by the partly redundant functions of a heterotrimeric kinesin-II and the homodimeric OSM-3 kinesin motor, whereas the distal segments are assembled by OSM-3 alone ^54,55^ (Figure 1A). We previously described the *osm-3(oy156)* temperature-sensitive (*ts*) allele which results in rapid truncation of channel cilia distal segments when adult *C. elegans* are shifted to the restrictive temperature of 30°C ^56^ (Figure 1A). We used this *ts* allele to test whether cilia can regenerate in the adult following truncation.

**Figure 1.**
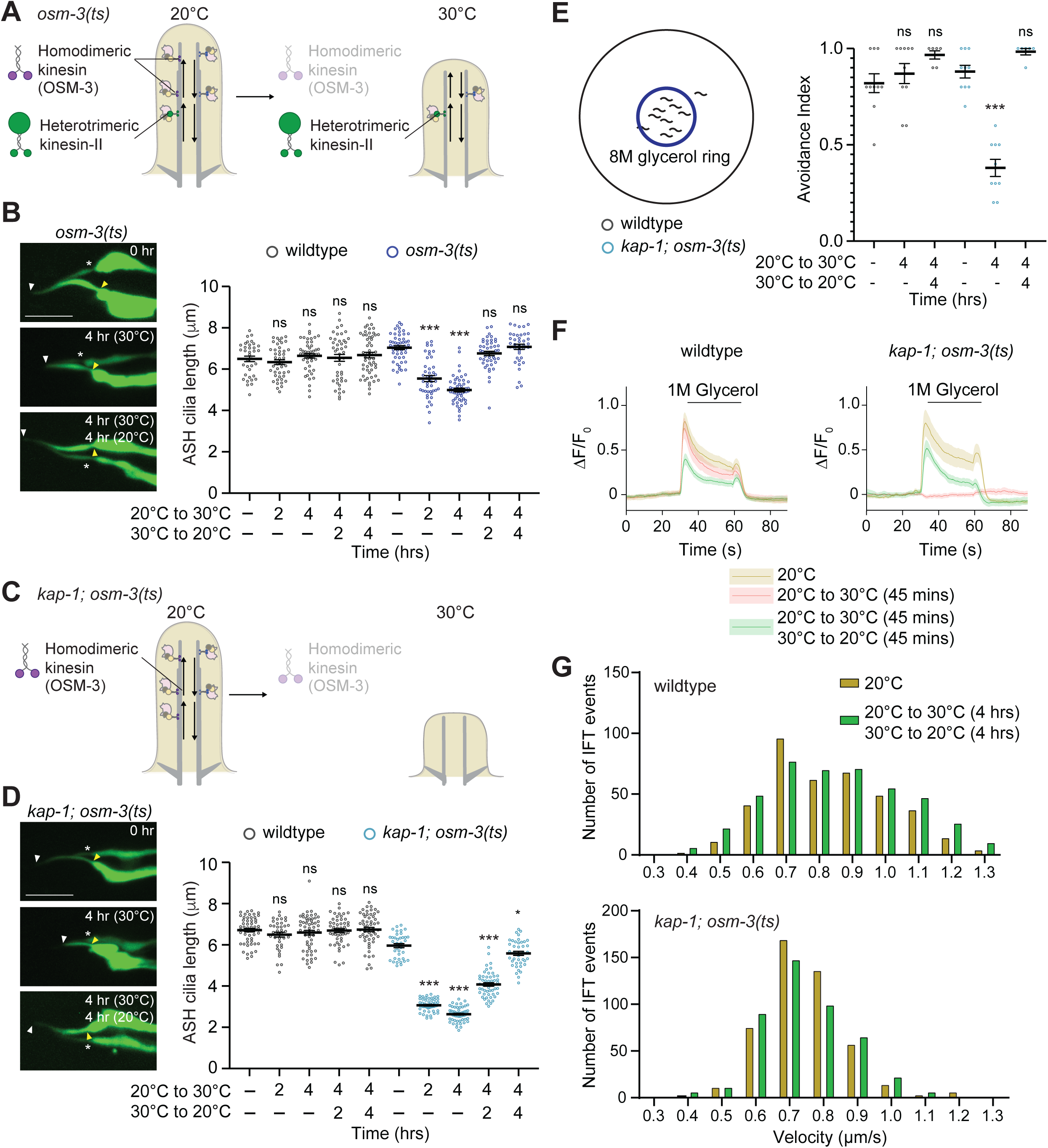
ASH cilia functionally regenerate upon truncation in adult *C. elegans*. **A,C)** Cartoons of ASH cilia in *osm-3(oy156ts)* (A) and *kap-1(ok676); osm-3(oy156ts)* (C) mutants before (left) and after (right) shift from the permissive (20°C) to the restrictive (30°C) temperature for 4+ hrs. Arrows indicate anterograde and retrograde IFT movement along the axoneme. **B,D)** (Left) Representative images of ASH cilia in *osm-3(oy156ts)* (B) and *kap-1(ok676); osm-3(oy156ts)* (D) mutants expressing *gfp* under the *sra-6* promoter at the indicated temperature shift conditions. Yellow/white arrowheads: cilia base/cilia tip; white asterisk: neighboring ASI cilium. Scale bars: 5 μm. (Right) Quantification of ASH cilia length in wildtype (black) and indicated mutants (dark and light blue) at the indicated temperature shift conditions. Each circle is the length of a single ASH cilium. * and ***: different at *P*<0.05 and 0.001, respectively, from unshifted within each genotype; ns: not significant (one-way ANOVA with Tukey’s multiple comparisons test). Errors are SEM. n≥36 each; 3 independent experiments. **E)** (Left) Cartoon of 8M glycerol avoidance assay. (Right) Quantification of 8M glycerol avoidance for wildtype (black) and *kap-1(ok676); osm-3(oy156ts)* mutants (blue) at the indicated temperature shift conditions. 0 and 1.0 indicate 0% and 100% avoidance, respectively. Each circle is a single assay of 10 animals. ***: different at *P*<0.001 from unshifted within each genotype; ns: not significant (one-way ANOVA with Tukey’s multiple comparisons test). Errors are SEM. n≥6 assays; 5 independent experiments. **F)** Mean GCaMP3 fluorescence changes in ASH to a 30s pulse of 1M glycerol in wildtype (left) and *kap-1(ok676); osm-3(oy156ts)* mutants (right) in the indicated temperature shift conditions. Shaded regions are SEM. n≥15 neurons each; 4 independent experiments. **G)** Histogram of velocities of reconstituted endogenously tagged OSM-6::split-GFP in ASH cilia in wildtype (top) and *kap-1(ok676); osm-3(oy156ts)* (bottom) animals. n≥19 animals per genotype; 3 independent experiments. Also see Figure S1.

We focused our analysis primarily on the monociliated ASH nociceptive sensory neurons as a representative of channel cilia. Cilia were visualized via expression of GFP under the *sra-6* promoter which drives expression strongly in ASH and more weakly in the ASI monociliated channel chemosensory neurons ^50,51,57^ (see Methods). Consistent with previous observations ^56^, shifting *osm-3(ts)* adult animals to 30°C resulted in loss of the distal segments of the rod-like cilia of ASH neurons (Figure 1B). Truncation was gradual and was complete by 4 hrs (Figure 1B). When animals were returned to the permissive temperature of 20°C, the ASH cilia regrew to their original length within 2 hrs (Figure 1B). Cilia length in wildtype ASH neurons was unaffected by these temperature shifts (Figure 1B). Both truncation and recovery of cilia morphology were markedly more rapid than that observed previously using the *che-3(nx159ts)* allele in the tail phasmid organ sensory neurons ^41^.

While only distal segments are truncated in *osm-3* mutants, both middle and distal segments are lost in *kap-1; osm-3* loss-of-function double mutants that lack both anterograde IFT motors ^54,55^ (Figure 1C). To determine whether both middle and distal segments can regrow, we performed similar temperature shift experiments in *kap-1; osm-3(ts)* animals. ASH cilia were shorter by 1 hr, and severely truncated by 4 hrs, after the temperature upshift (Figure 1D, Figure S1). Upon returning to the permissive temperature, significant elongation of these cilia was observed within 1 hr and these cilia regrew to nearly their original length by 4 hrs (Figure 1D, Figure S1). We infer that both the middle and distal segments of ASH cilia regrow rapidly following acute truncation in adult *C. elegans*.

We next assessed whether sensory functions are restored in neurons with regrown cilia. ASH detects and drives avoidance of high osmolarity ^58^. Wildtype animals placed within a ring of 8M glycerol remain within the ring since they avoid this chemical. Animals lacking ASH cilia rapidly escape the ring since they fail to avoid high osmolarity (*osm* or osmotic avoidance defective) ^59^. At 20°C, *kap-1; osm-3(ts)* mutants avoided glycerol (Figure 1E). However, upon shifting to 30°C, these animals escaped the ring as ASH cilia were truncated (Figure 1E). Glycerol avoidance was fully restored after returning to 20°C correlating with cilia regrowth (Figure 1E). Consistently, glycerol evoked a robust increase in intracellular calcium in ASH neurons in *kap-1; osm-3(ts)* double mutants at the permissive temperature, but this response was abolished at the restrictive temperature (Figure 1F). Calcium responses were restored upon return to the permissive temperature (Figure 1F). Calcium dynamics were unaltered in wildtype animals at any examined temperature condition (Figure 1F). We conclude that recovery of cilia length restores neuronal functions.

We also determined whether IFT dynamics are restored upon cilia regrowth by analyzing the movement of the IFT-B particle component OSM-6/IFT52. To visualize IFT in ASH we used the previously described *osm-6(oy166)* allele in which *gfp_11_* sequences are inserted at the endogenous *osm-6* locus via gene editing ^56^. GFP was reconstituted in ASH via expression of the GFP_1-10_ fragment under the *sra-6* promoter ^56,57^. Movement of OSM-6::split-GFP in regrown ASH cilia was comparable to that observed in *kap-1; osm-3(ts)* animals at the permissive temperature (Figure 1G). Together, these results indicate that both sensory functions and IFT are restored in regrown ASH cilia. Henceforth we refer to these structures as regenerated cilia.

### The DAF-19 RFX transcription factor is necessary for cilia regeneration but not maintenance in the adult

The conserved DAF-19 RFX transcription factor is the master ciliogenic gene in *C. elegans* ^43^. This protein regulates the expression of all core ciliary genes including IFT genes by directly binding X box motifs in their upstream regulatory regions ^60–62^. Consequently, cilia fail to form in *daf-19* mutants ^43^. Although transcriptomic studies have suggested that DAF-19 is partly dispensable for cilia maintenance in later postembryonic stages ^63^, a role for this transcription factor in maintaining and regenerating cilia in the adult has not been tested.

To determine whether DAF-19 is required for cilia regeneration, we conditionally depleted DAF-19 specifically in the adult in ASH/ASI neurons. We generated an endogenously tagged *daf-19* allele fused to an auxin-inducible degron (AID) ^64,65^ (Figure 2A). Animals carrying this allele did not exhibit cilia defects as assessed via their normal cilia lengths and ability to uptake a lipophilic dye (Figure 2B, Figure S2A), indicating that this allele retains function. We first determined whether acute depletion of DAF-19 in the adult disrupts cilia. Adult animals expressing *daf-19::AID* with or without the auxin receptor TIR1 expressed under the *sra-6* promoter in ASH/ASI (Figure 2A) were transferred to plates containing either the ethanol control or 4 mM auxin, and cilia morphology was assessed after 8 or 20 hrs (Figure 2B). ASH cilia length was either slightly increased or unaffected under all examined conditions (Figure 2B). To confirm effective depletion, we examined GFP levels in animals expressing an endogenous functional *daf-19::AID::gfp* allele (Figure S2A) together with TIR1 expressed in ASH/ASI on plates containing auxin. GFP levels in both ASH and ASI appeared to be markedly reduced by 30 mins and were undetectable by 4 hrs of auxin exposure (Figure S2B). We conclude that DAF-19 is not necessary to maintain cilia in the adult.

**Figure 2.**
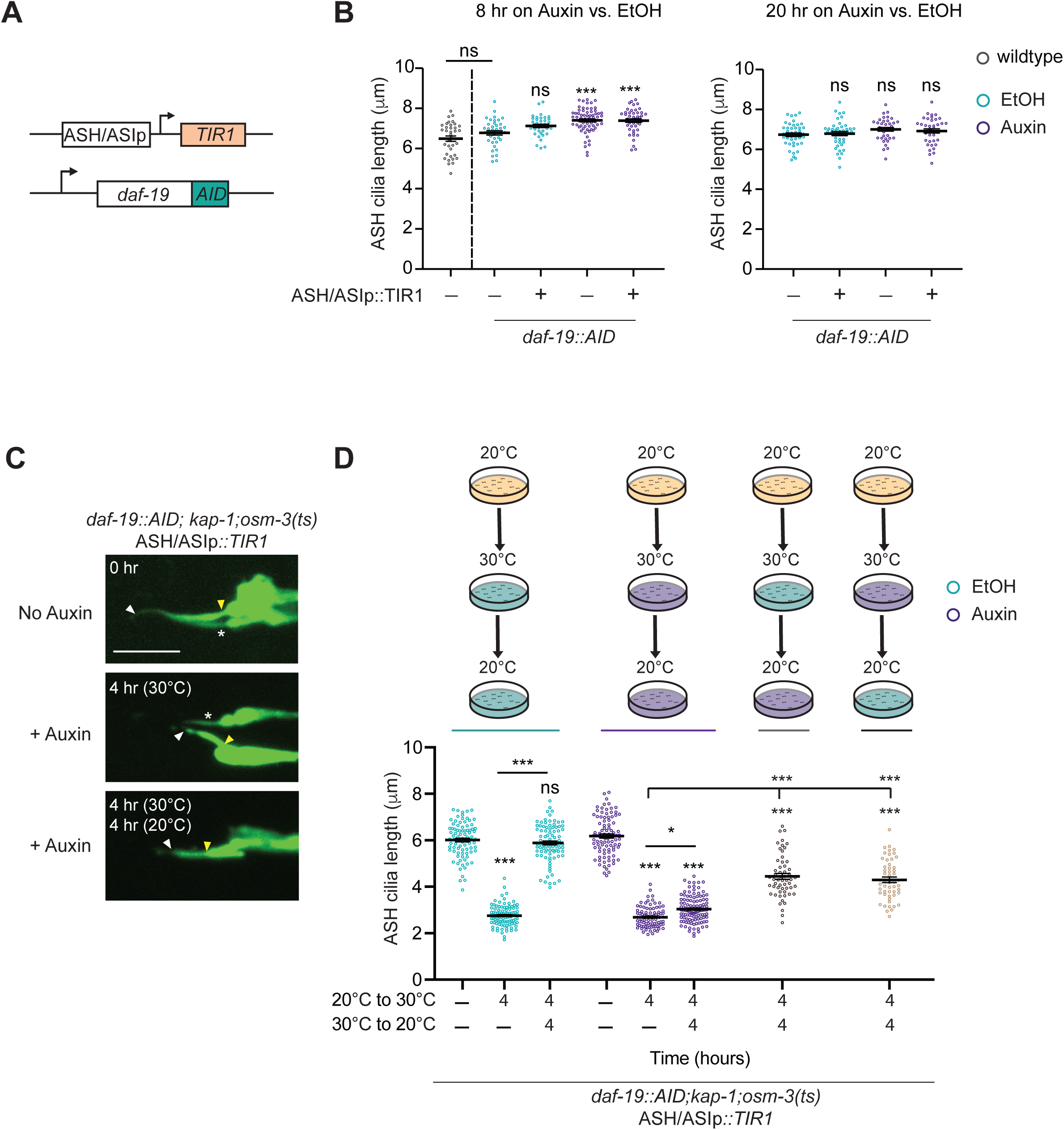
DAF-19 is required for cilia regeneration but not cilia maintenance. **A)** Schematic of endogenously tagged *daf-19::AID* together with TIR1 expressed under the *sra-6* promoter from an extrachromosomal array. The *sra-6* promoter used drives expression in ASH and ASI in the head ^57^. **B)** Quantification of ASH cilia length in animals containing *daf-19::AID* with or without the *sra-6*p*::TIR1* transgene on ethanol control (blue circles) or auxin-containing (purple circles) plates for 8 (left) or 20 (right) hrs. ASH cilia length in wildtype animals under standard growth conditions (black circles) is shown for comparison on the left (data are repeated from Figure 1B). Each circle is the length of a single ASH cilium. ***: different from ethanol control without TIR1 expression at *P*<0.001; ns: not significant (unpaired Welch’s t-test). n≥37 each; 3 independent experiments. **C)** Representative images of ASH cilia expressing GFP in *daf-19(oy204); kap-1(ok676); osm-3(oy156ts)* mutants together with *sra-6*p*::TIR1* in the indicated temperature shift conditions with or without auxin. Yellow/white arrowheads: cilia base/cilia tip; white asterisk: neighboring ASI cilium. Scale bar: 5 μm. **D)** (Top) Schematic of experimental conditions associated with cilia length quantifications (bottom). Each circle is the length of a single ASH cilium. * and ***: different at *P*<0.05 and 0.001, respectively from unshifted within each condition or between indicated; ns: not significant (one-way ANOVA with Tukey multiple comparison’s test or unpaired Welch’s t-test. Errors are SEM. n≥50 each; 6 independent experiments for continuous auxin or EtOH treatment, and 3 independent experiments for auxin treatment only during truncation or regrowth. Also see Figure S2.

We next asked whether DAF-19 is necessary for cilia truncation and/or regeneration by shifting adult *kap-1; osm-3(ts)* mutants carrying the *daf-19::AID* allele and expressing TIR1 in ASH/ASI to either the restrictive or permissive temperature in the continuous presence of ethanol or auxin and then assessing cilia length. Cilia truncated robustly upon a shift to 30°C in both conditions (Figure 2C, 2D, Figure S2C). However, while animals on ethanol regenerated their cilia upon return to 20°C, cilia failed to elongate in animals continuously exposed to auxin, indicating a requirement for DAF-19 in cilia regrowth (Figure 2C, 2D). ASH cilia also exhibited significant regrowth defects in animals placed on auxin only during the upshift to 30°C or only during the downshift to 20°C, although we observed partial regrowth under both conditions (Figure 2D). We note that DAF-19::GFP levels were not fully restored and cilia did not fully regenerate even at >20 hrs following auxin removal (Figure S2B, S2C) indicating that we were unable to deplete DAF-19 specifically during cilia truncation. Together, these results suggest that while DAF-19 is dispensable for maintenance of channel cilia in the adult, this transcription factor is essential for cilia regrowth, and may function both during cilia truncation and regeneration phases to promote efficient regeneration.

### The ciliogenic DAF-19d isoform regulates cilia regeneration

The *daf-19* locus encodes multiple protein isoforms which share common C-terminal sequences but differ at their N-termini ^63,66–68^ (Figure 3A). While the *daf-19d* isoform (previously *daf-19c*) has been specifically implicated in ciliogenesis ^63,66^, *daf-19a/b* are broadly expressed and regulate synaptic functions ^63,66^, *daf-19c* (previously *daf-19m*) expression is restricted to the IL2 sensory neurons in hermaphrodites ^68^, and the function of *daf-19e* is unknown. Since we were unable to selectively mutate *daf-19d* without also affecting other isoforms, we first examined animals carrying the *daf-19(tm5562)* allele to determine whether *daf-19d* regulates cilia regeneration. The *daf-19(tm5562)* mutation disrupts the longer *daf-19a* and *daf-19b* isoforms and these mutants exhibit phenotypes consistent with loss of these isoforms while retaining *daf-19d* function ^63^ (Figure 3A). We observed no defects in either cilia truncation or regrowth in *daf-19(tm5562); kap-1; osm-3(ts)* triple mutants (Figure 3B), supporting the hypothesis *daf-19a/b* does not regulate cilia regeneration. To further assess a requirement for *daf-19d* in cilia regeneration, we expressed a genomic fragment that includes the upstream regulatory and coding sequences of *daf-19d* ^66^ (Figure 3A) in *kap-1; osm-3(ts)* animals carrying the endogenously tagged *daf-19::AID* allele and TIR1 expressed in ASH/ASI, and compared cilia lengths on auxin. While ASH cilia failed to regenerate at the permissive temperature in the control strain in the presence of auxin, expression of *daf-19d* was sufficient to significantly rescue cilia regrowth (Figure 3C). We conclude that *daf-19d* regulates cilia regeneration in adult *C. elegans* although we are unable to formally exclude a possible role of *daf-19c* and/or *daf-19e*.

**Figure 3.**
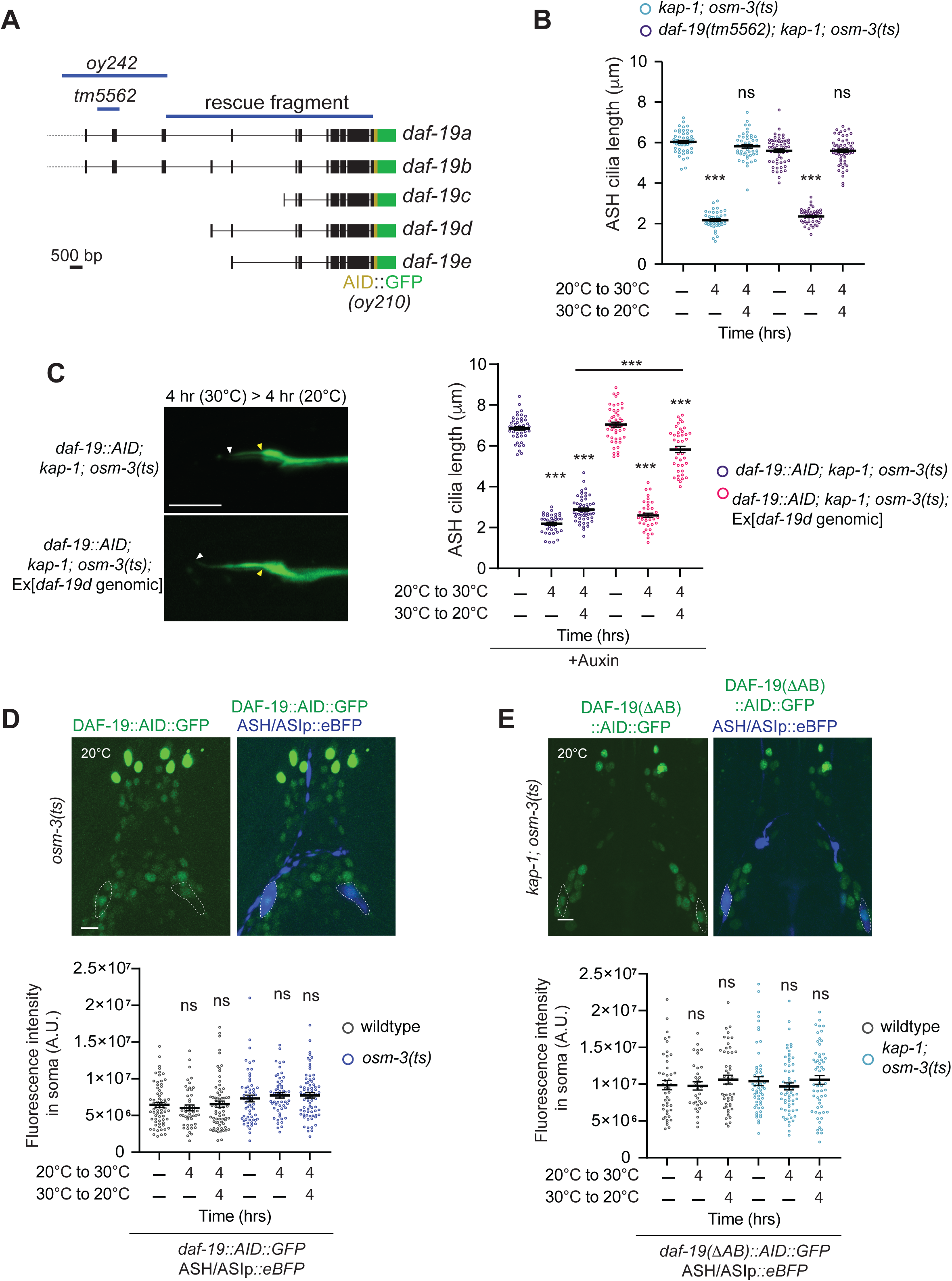
The ciliogenic *daf-19d* isoform regulates cilia regeneration. **A)** Schematic showing predicted *daf-19* isoforms (from www.wormbase.org) endogenously tagged with *AID::gfp* sequences. The molecular lesions in *daf-19(tm5562)* and *daf-19(oy242)* and the genomic sequences included in the *daf-19d* rescue construct are indicated. **B)** Quantification of ASH cilia length in *kap-1; osm-3(ts)* or *daf-19(tm5562); kap-1; osm-3(ts)* mutants at the indicated temperature shift conditions. Each circle is the length of a single ASH cilium. ***: different at *P*<0.001 from unshifted within each genotype; ns: not significant (one-way ANOVA with Tukey multiple comparison’s test). Errors are SEM. n≥40 each; 3 independent experiments. **C)** (Left) Representative images of ASH cilia length in animals of the indicated genotypes at the shown temperature shift conditions. Anterior is at left. Scale bar: 5 μm. White and yellow arrowheads indicate cilia tip and base, respectively. (Right) Quantification of ASH cilia length in animals of the indicated genotypes at the shown temperature shift conditions. Each circle is the length of a single ASH cilium. ***: different at *P*<0.001 from unshifted within each genotype or between indicated; ns: not significant (one-way ANOVA with Tukey multiple comparison’s test or unpaired Welch’s t-test). Errors are SEM. n≥37 each; 3 independent experiments. **D,E)** (Top) Representative images of reporter expression from endogenously tagged *daf-19(oy210)* (D) and *daf-19(oy210 oy242Δa/b)* (E) alleles in ASH/ASI soma (dashed outlines) marked via expression of *sra-6*p*::eBFP* in the indicated genetic backgrounds and temperature shift conditions. Anterior at top. Scale bars: 5 μm. (Bottom) Quantification of reporter fluorescence intensities in ASH/ASI soma of endogenously reporter tagged *daf-19(oy210)* (D) and endogenously reporter-tagged *daf-19(oy210 oy242Δa/b)* (E) in animals of the indicated genotypes in the shown temperature shift conditions. Each circle is the value from an ASH and/or ASI soma in one amphid organ. ns: not significant from unshifted within each genotype (one-way ANOVA with Tukey multiple comparison’s test). Errors are SEM. n≥34 each; 2 independent experiments. Also see Figure S2.

We next examined whether DAF-19 protein levels are altered during cilia truncation and regeneration in adults. We did not detect altered expression of an endogenous reporter-tagged DAF-19 protein (Figure 3A) in ASH/ASI soma regardless of cilia length (Figure 3D). Since this allele contains *gfp* sequences fused to the common C-terminus (Figure 3A), we reasoned that upregulation of *daf-19d* alone may be masked by other isoforms. To examine *daf-19d* levels, we generated the *daf-19(oy210 oy242)* allele designed to disrupt expression of *daf-19a* and *daf-19b* while retaining expression of *daf-19c*, *daf-19d* and *daf-19e* isoforms (Figure 3A). *daf-19(oy210 oy242Δa/b)* mutants did not exhibit dye-filling defects indicating that this allele does not disrupt ciliogenesis (Figure S2A). Reporter expression driven from the *daf-19(oy210 oy242Δab)* allele was also unaltered upon cilia truncation and regrowth (Figure 3E, Figure S2D). These observations suggest that although DAF-19d is required for cilia regeneration, its activity during this process is likely regulated by mechanisms other than altered expression (see Discussion).

### DAF-19 regulates distinct temporal dynamics of IFT gene expression during cilia truncation and regrowth

Complete regrowth of *Chlamydomonas* flagella following severing requires transcriptional upregulation of IFT and flagellar building block genes ^69–75^. Given the requirement for *daf-19d* in cilia regeneration, we asked whether IFT genes are also upregulated during cilia regeneration in *C. elegans*, and whether this upregulation is mediated by DAF-19.

To examine IFT gene expression during cilia truncation and regrowth, we generated strains in which IFT genes are tagged at their endogenous loci with either *SL2::gfp_11_* or *gfp_11_* sequences alone, together with expression of *gfp_1-10_* under the *sra-6* promoter to visualize reconstituted GFP selectively in ASH/ASI (Figure 4A, Figure S3A). Insertion of the *SL2* trans-splice leader sequence allows monitoring of transcriptional changes independent of protein localization. We examined the *osm-6/IFT52, osm-5/IFT88*, and *osm-1/IFT172* genes which encode conserved components of the IFT-B complex ^76,77^. Loss-of-function mutations in each of these genes result in severe ciliary structural defects ^78,79^.

**Figure 4.**
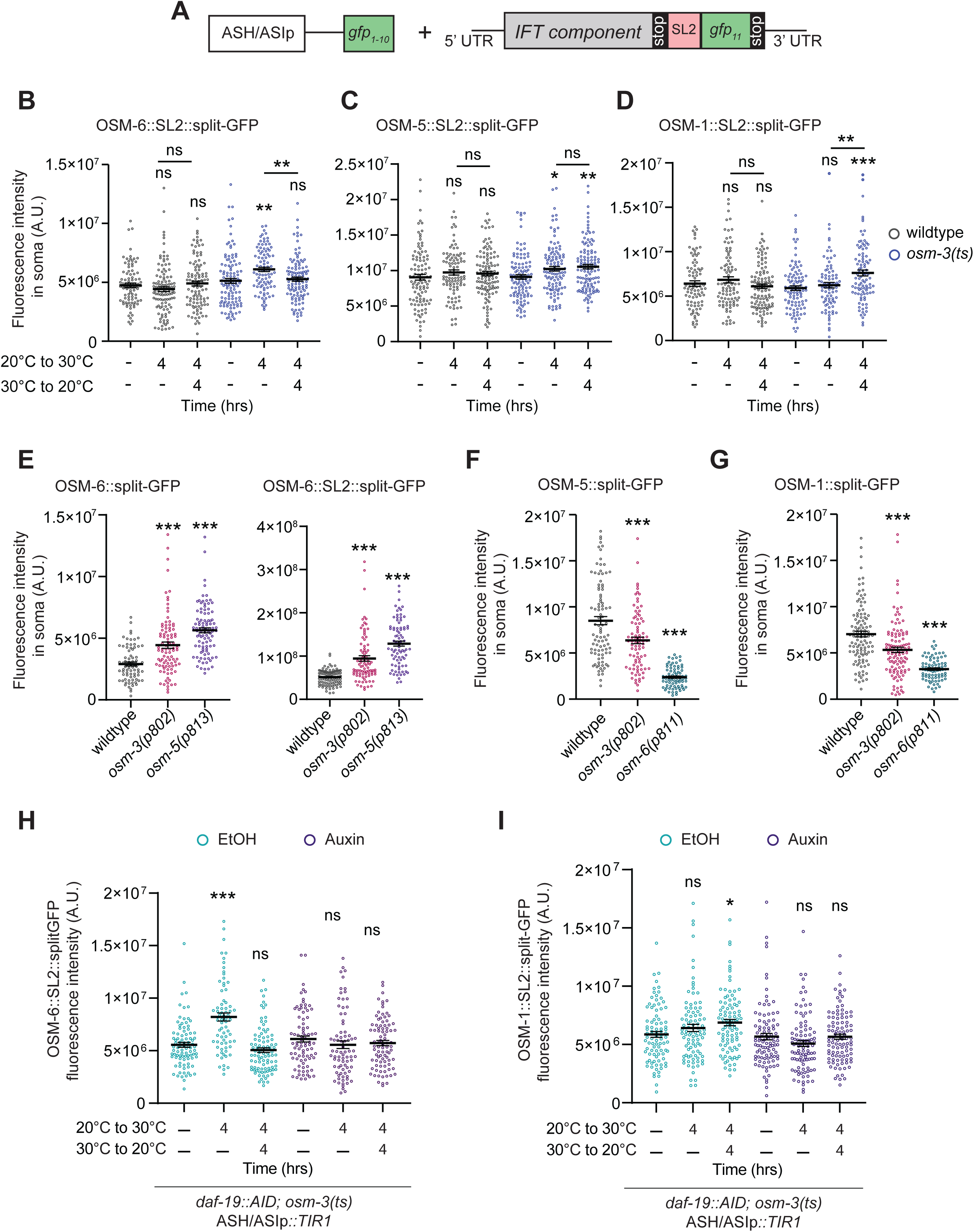
DAF-19 regulates temporal expression dynamics of IFT genes during cilia truncation and regeneration. **A)** Schematic of endogenously tagged transcriptional reporter expression. **B-D)** Quantification of fluorescence intensities in ASH/ASI soma of wildtype (black) or *osm-3(oy156ts)* (blue) animals expressing the indicated IFT transcriptional reporters in the shown temperature shift conditions. Each circle is the value from an ASH and/or ASI soma in one amphid organ. *, ** and ***: different at *P*<0.05, 0.01 and 0.001, respectively, from unshifted within each genotype or between indicated; ns: not significant (one-way ANOVA with Tukey multiple comparison’s test or unpaired Welch’s t-test). Errors are SEM. n≥85 each; 3 independent experiments. **E-G)** Quantification of fluorescence intensities of the indicated reporter-tagged IFT proteins or transcriptional reporter in the ASH/ASI soma of wildtype or IFT gene mutants. Each circle is the Each circle is the value from an ASH and/or ASI soma in one amphid organ. ***: different at *P*<0.001 from wildtype (one-way ANOVA with Tukey’s multiple comparison test). Errors are SEM. n≥74 each; 3 independent experiments. **H,I)** Quantification of endogenously tagged OSM-6::SL2::split-GFP (H) or OSM-1::SL2::split-GFP (I) fluorescence intensities in ASH/ASI soma of *daf-19(AID); osm-3(ts)* mutants expressing *sra-6*p*::TIR1* in the indicated conditions. Each circle is the value from an ASH and/or ASI soma in one amphid organ. * and ***: different at *P*<0.05 and 0.001, respectively, from unshifted within each condition; ns: not significant (one-way ANOVA with Tukey’s multiple comparison test). Errors are SEM. n≥73 each; 3 independent experiments. Also see Figure S3.

Expression of *osm-6::SL2::split-GFP* was significantly upregulated in ASH/ASI in *osm-3(ts)* mutants when the distal segments were fully truncated, and subsequently returned to baseline levels as cilia regenerated (Figure 4B). OSM-6::split-GFP protein levels in ASH/ASI soma showed a similar pattern of upregulation although protein levels in cilia appeared unaltered (Figure S3B). In contrast, *osm-5::SL2::split-GFP* was also upregulated upon cilia truncation but was then subsequently maintained upon regeneration (Figure 4C), although no changes in OSM-5 protein levels were detected either in ASH/ASI soma or cilia (Figure S3C). *osm-1* exhibited distinct expression dynamics such that *osm-1* expression was significantly upregulated only upon cilia regrowth with the protein exhibiting similar expression changes in the ASH/ASI soma but not in cilia (Figure 4D, Figure S3D). Together, these results indicate that a subset of IFT genes is transcriptionally upregulated during cilia truncation and regeneration with distinct temporal patterns of expression.

We next asked whether altered IFT gene expression is correlated with cilium length and/or reflects a distinct cellular state associated with acute cilium disassembly or regeneration in the adult. To distinguish between these possibilities, we examined IFT gene expression in animals carrying loss-of-function mutations in the *osm-3* kinesin motor or other IFT genes which result in truncated cilia throughout development. As observed during acute conditional cilia truncation, OSM-6 protein levels and expression of the *osm-6* transcriptional reporter were elevated in ASH/ASI soma in both *osm-3* and *osm-5* loss-of-function mutants (Figure 4E). In contrast, levels of both OSM-5 and OSM-1 proteins were instead markedly reduced in animals carrying loss-of-function alleles of either *osm-3* or *osm-6* (Figure 4F, 4G). In all cases, expression changes were more pronounced in *osm-5* or *osm-6* mutants that exhibit severely truncated cilia than in *osm-3* mutants in which only the distal ciliary segments fail to assemble (Figure 4E-G). These observations suggest that while induction of *osm-6* expression may partly scale with cilia length regardless of condition, the expression of other IFT genes is regulated in a more complex manner by both cilia and developmental state.

We tested whether DAF-19 is required for the upregulation of IFT genes during cilia truncation and regeneration. Consistent with the observation that DAF-19 is dispensable for cilia maintenance in the adult, expression levels of IFT genes were unaffected upon depletion of DAF-19 under control growth conditions (Figure 4H, 4I). However, the truncation and regeneration-associated upregulation of both *osm-6::SL2::gfp* and *osm-1::SL2::gfp* was abolished upon depletion of DAF-19 in the adult (Figure 4H, 4I). We conclude that while DAF-19 is not required to maintain IFT gene expression in the adult, this transcription factor is necessary for transcriptional upregulation of IFT gene expression likely supporting cilia regeneration.

### The DLK-1 MAPKKK, and DAF-19-mediated induction of the CEBP-1 C/EBP transcription factor, promote cilia regeneration

Both axons and cilia are microtubule-based structures with their plus-ends oriented distally. Peripheral axons exhibit robust regeneration following injury in adults, and this regeneration is mediated in part via injury-induced changes in gene expression ^28,80,81^. We tested whether molecules implicated in axon regeneration also play a role in regrowth of sensory cilia. The DLK dual leucine-zipper kinase is activated by disruption of the microtubule cytoskeleton upon axon injury and promotes axon regeneration via downstream transcriptional programs ^46,47,82–87^. Recent work in *C. elegans* has also shown that mutations in IFT genes increase DLK-1 accumulation in sensory cilia and that this kinase acts via the CEBP-1 C/EBP transcription factor to protect a subset of sensory cilia from degeneration ^88,89^. We investigated whether the DLK-1 signaling pathway contributes to efficient cilia regrowth.

Loss-of-function mutations in *dlk-1* caused only minor defects in ASH cilia length (Figure S4A). We also did not observe effects on ASH cilia truncation in *dlk-1; kap-1; osm-3(ts)* triple mutants upon an upshift to 30°C (Figure 5A). However, upon return to the permissive temperature of 20°C, loss of *dlk-1* resulted in a failure of cilia to fully regenerate to their original length by 4 hrs (Figure 5A). This defect was rescued upon expression of wildtype *dlk-1* sequences in ASH/ASI (Figure 5A). The partial regeneration of cilia in *dlk-1* mutants after 4 hrs at the permissive temperature suggested that loss of *dlk-1* may delay but not fully abolish cilia regeneration. Consistent with this notion, cilia in *dlk-1* mutants eventually regenerated to the wildtype length after 16 hrs at the permissive temperature (Figure S4B). Previous work reported that movement of the kinesin-II motor is slowed in ciliary middle segments in *dlk-1* mutants ^89^. However, we did not detect changes in IFT-mediated transport of endogenously tagged OSM-6 in *dlk-1* mutants (Figure S4C), suggesting that slower IFT dynamics may not be the primary cause of delayed regrowth in *dlk-1* mutants. We infer that DLK-1 is required for efficient cilia regeneration.

**Figure 5.**
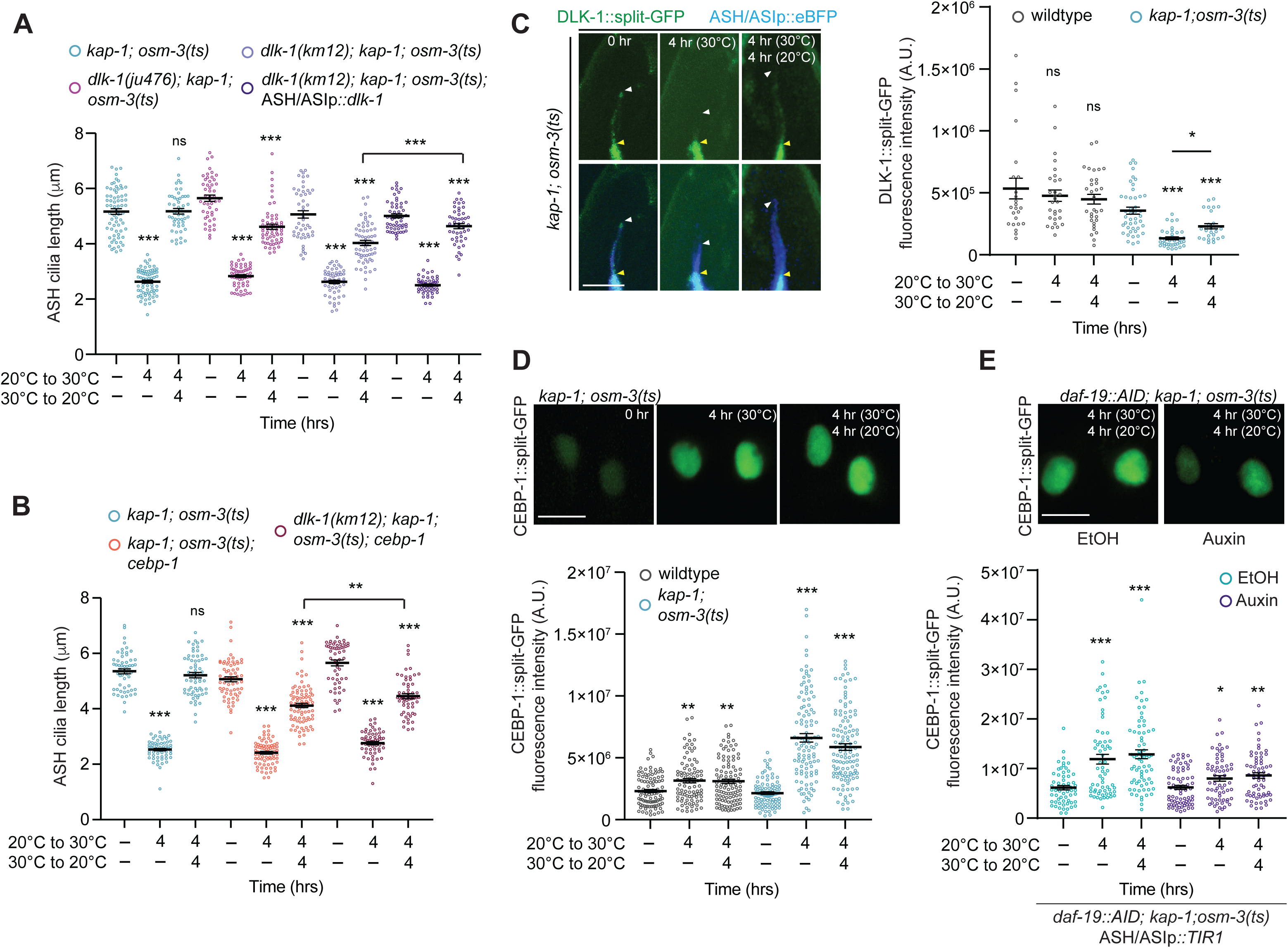
DLK-1 and DAF-19-regulated CEBP-1 induction promote cilia regeneration. **A,B)** Quantification of ASH cilia length in the shown genetic backgrounds at the indicated temperature shift conditions. Wildtype *dlk-1* sequences were expressed in ASH/ASI under the *sra-6* promoter. Each circle is the length of a single ASH cilium. **, ***: different at *P*<0.01 and 0.001, respectively, from unshifted within each genotype or between indicated; ns: not significant (one-way ANOVA with Tukey’s multiple comparison test or unpaired Welch’s t-test). Errors are SEM. n≥48 each; >3 independent experiments. **C)** (Left) Representative images of endogenously tagged DLK-1:split-GFP expression in ASH cilia at the indicated temperature shift conditions and genetic backgrounds. Yellow/white arrowheads: cilia base/cilia tip. Anterior is at top. Scale bar: 5 μm. (Right) Quantification of fluorescence intensities of DLK-1::split-GFP in ASH cilia in the indicated genetic backgrounds and temperature shift conditions. Each circle is the value from a single ASH cilium. * and ***: different at *P*<0.05, 0.01 and 0.001, respectively, from unshifted within each genotype or between indicated; ns: not significant (one-way ANOVA with Tukey’s multiple comparison test or unpaired Welch’s t-test). Errors are SEM. n≥24 each; 3 independent experiments. **D,E)** (Top) Representative images of endogenously tagged CEBP-1::split-GFP in ASH/ASI soma at the indicated temperature shift conditions and in ethanol control or auxin (E). Scale bars: 5 μm. (Bottom) Quantification of fluorescence intensities of CEBP-1::split-GFP in ASH/ASI soma in the indicated genetic backgrounds, temperature shift conditions, and in ethanol control or auxin (E). Each circle is the value from an ASH and/or ASI soma in one amphid organ. *, ** and ***: different at *P*<0.05, 0.01 and 0.001, respectively, from unshifted within each genotype; ns: not significant (one-way ANOVA with Tukey’s multiple comparison test). Errors are SEM. n>60 each; 3 independent experiments. Also see Figure S4.

DLK-1 regulates axon regeneration through well-characterized MAP kinase signaling cascades (Figure S4D) ^29^. To determine whether these pathways contribute to cilia regeneration, we examined mutants in downstream signaling components. Mutations in *mlk-1* MAPKKK or *pmk-3* MAPK mutants did not affect cilia regrowth, although loss of *kgb-1* JNK-like MAPK caused a significant defect in cilia regeneration (Figure S4E). We also examined whether the downstream transcription factor CEBP-1 is required for cilia regeneration. As observed for *dlk-1* mutants, loss of *cebp-1* had only a minor effect on ASH cilia length (Figure S4A) but affected ASH cilia regeneration in the *kap-1; osm-3(ts)* mutant background (Figure 5B). This regeneration defect was altered further to only a minor extent in *dlk-1; cebp-1* double mutants suggesting that these molecules act in the same pathway to promote cilia regeneration (Figure 5B).

DLK-1 levels are low in cilia in wildtype animals but are increased upon cilia truncation in IFT mutants ^88^. Similarly, CEBP-1 levels increase in the soma of sensory neurons in IFT mutants ^88^. Endogenously tagged DLK-1::split-GFP ^88^ was present at low levels in ASH/ASI cilia in both wild-type animals and in *kap-1; osm-3(ts)* mutants at 20°C (Figure 5C). DLK-1 levels were further decreased in these cilia upon truncation in the double mutant, but increased significantly although not to wild-type levels upon cilia regrowth (Figure 5C). In contrast, levels of a nuclear-localized endogenously tagged CEBP-1::split-GFP reporter ^90^ were significantly increased in ASH/ASI soma of *kap-1; osm-3(ts)* mutants upon truncation and remained elevated when cilia were fully regrown (Figure 5D, 5E). Auxin-mediated depletion of DAF-19 abolished *cebp-1* induction upon cilia truncation and regrowth (Figure 5E). Together, these results indicate that CEBP-1 and DLK-1 promote efficient cilia regeneration, and that in addition to upregulating IFT genes, DAF-19 also induces *cebp-1* expression in truncated cilia.

### Glial signals differentially influence neuron type-specific dynamics of cilia truncation and regeneration

Sensory cilia in *C. elegans* exhibit remarkably diverse morphologies ^50,51,78^. We asked whether distinct cilia types can undergo truncation and regeneration in the adult. Unlike the monociliated ASH or ASI neurons, the ADF neurons contain two cilia that are also present in the amphid channel (Figure 6A, 6B) ^50,51,78^. In *kap-1; osm-3(ts)* mutants, these cilia truncated and regrew with dynamics similar to those observed for ASH cilia following shifts to the restrictive and permissive temperatures, respectively (Figure 6B). The AWB olfactory neurons contain morphologically complex cilia that are embedded in the amphid sheath glial processes unlike channel cilia (Figure 6A, 6C) ^50,51,78^. In *kap-1; osm-3(ts)* mutants, AWB cilia truncated and regenerated, but did so on significantly longer timescales than ASH cilia. AWB cilia were only partly truncated even after 24 hrs at 30°C (average AWB cilium length in *kap-1; osm-3(lof)* and *kap-1; osm-3(ts)* double mutants is 2.9 ± 0.2 μm and 6.4 ± 2.1 μm, respectively), and these cilia failed to regrow to their original length even after 8 hrs at 20°C (Figure 6C). We infer that a subset of diverse sensory cilia types in adult *C. elegans* are able to truncate and regrow, but that the dynamics of these processes vary in a neuron type-specific manner.

**Figure 6.**
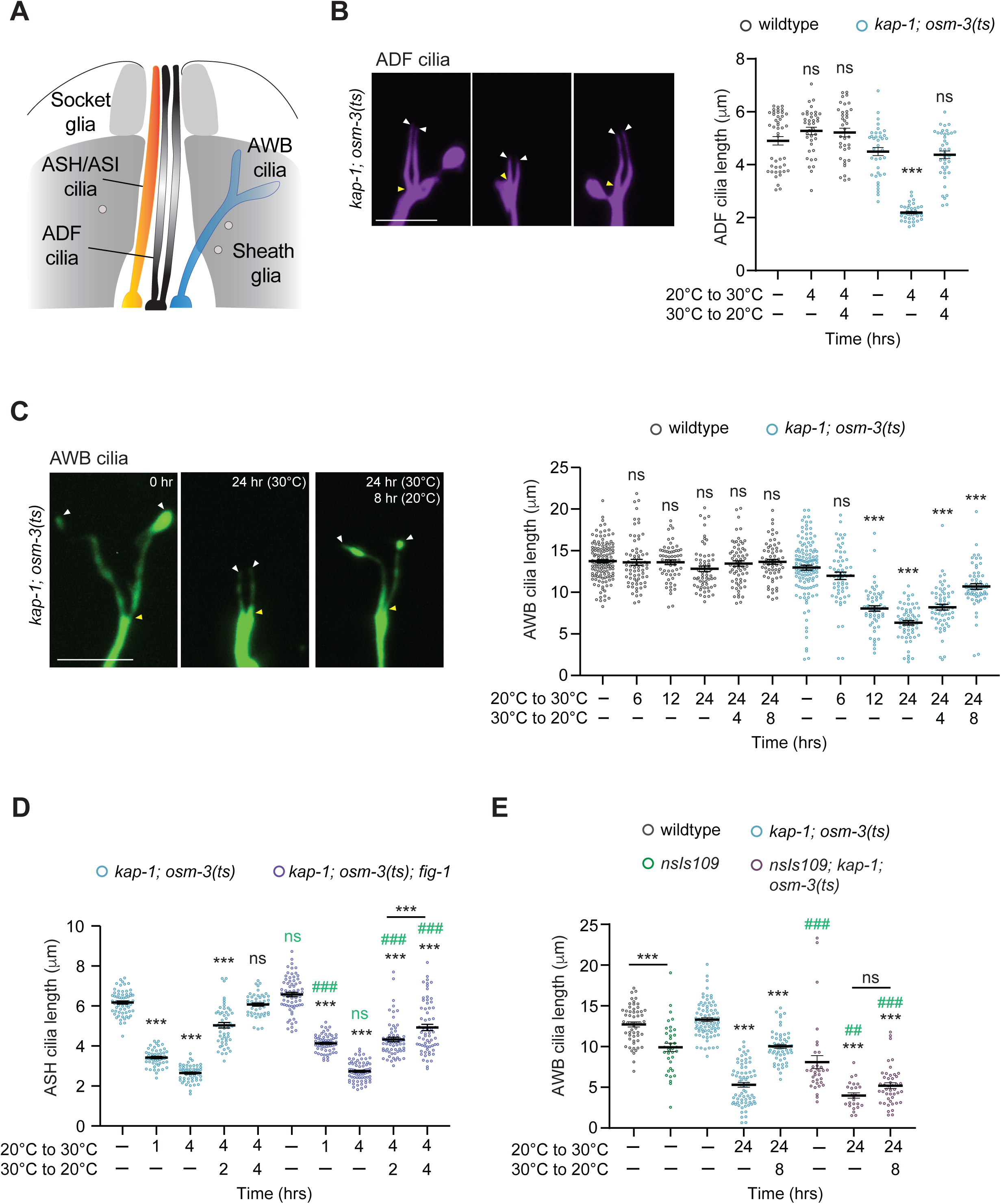
Glia modulate cilia regeneration dynamics in a cell type-specific manner. **A)** Schematic of the cilia of the ASH/ASI, ADF, and AWB neurons with associated sheath and socket glia in the amphid sense organ of the head. Anterior is at top. Adapted from ^78^. **B,C)** (Left) Representative images of ADF (B) and AWB (D) cilia lengths (right) in the shown genetic backgrounds at the indicated temperature shift conditions. Anterior at top. White/yellow arrowheads: cilia tip and base, respectively. Scale bars: 5 μm. (Right) Quantification of ADF (B) and AWB (D) cilia lengths in the shown genetic backgrounds at the indicated temperature shift conditions. Each circle is the average length of both ADF or AWB cilia from a single neuron. ***: different at *P*<0.001 from unshifted within each genotype; ns: not significant (one-way ANOVA with Tukey’s multiple comparison test). Errors are SEM. n≥38 each; >3 independent experiments. **D,E)** Quantification of ASH (D) and AWB (E) cilia length in the shown genetic backgrounds at the indicated temperature conditions. Each circle is the length of a single ASH cilium (D) or the average length of both AWB cilia from a single neuron (E). ***: different at *P*<0.001 from unshifted within each genotype or between indicated; ns (black): not significant (one-way ANOVA with Tukey’s multiple comparison test). ## and ###: different at *P*<0.01 and 0.001, respectively, from the *kap-1; osm-3(ts)* genotype at the corresponding timepoint; ns (green): not significant (unpaired Welch’s t-test). Errors are SEM. n≥25 each; >3 independent experiments. Also see Figure S5.

While mammalian peripheral axons regenerate robustly following axotomy, central axons fail to do so efficiently, in part due to inhibitory signals from glia and the extracellular matrix ^32,33^. In *C. elegans*, the amphid sheath glia regulate cilia morphology and protect cilia from degeneration ^91–94^. Disruption of channel cilia results in accumulation of glia-secreted extracellular matrix into the amphid channel; this glial response is neuroprotective ^92^. We tested whether glia influence neuron-specific dynamics of cilia truncation and regeneration.

We ablated the sheath glia using a previously described strain in which diptheria toxin A is expressed specifically in the sheath glia following amphid development ^91^. We noted significant defects in ASH cilia morphology in the *kap-1; osm-3(ts)* double mutant strain in the glia-ablated background even at the permissive temperature that precluded further analyses of cilia truncation and regrowth in this strain (Figure S5A). Channel cilia integrity has been proposed to be monitored by interactions between the cilia-localized DGS-1 transmembrane and glia-expressed thrombospondin domain FIG-1 proteins ^92^. Mutations in either *dgs-1* or *fig-1* trigger the glial protective secretory response even in animals with intact channel cilia ^92^.

Mutations in *fig-1* did not alter ASH cilia length at the permissive temperature (Figure 6D). However, in *kap-1; osm-3(ts); fig-1* mutants ASH cilia truncated more slowly upon a shift to 30°C, and failed recover to their original length after 4 hrs at the permissive temperature (Figure 6D). These observations suggest that the glial secretome may modulate the dynamics of channel cilia truncation and regrowth.

Although AWB cilia structure was reported to be largely retained upon sheath glia ablation ^91^, we found that AWB cilia length and morphology were also significantly affected in these animals and further shortened in a *kap-1; osm-3(ts)* background even at the permissive temperature (Figure 6E, Figure S5B). AWB cilia exhibited additional truncation upon a temperature upshift and failed to regrow upon return to the permissive temperature (Figure 6E, Figure S5B). These observations further support a role for glia in modulating cilia truncation and regrowth, and suggest that these mechanisms operate in a neuron type-specific manner in the adult.

## DISCUSSION

Here we show that sensory neuron cilia in adult *C. elegans* can regenerate and restore neuronal function. We find that the ciliogenic DAF-19 RFX transcription factor is not necessary for maintaining cilia in the adult but is essential for cilia regeneration. Cilia truncation and regrowth induce the expression of a subset of IFT genes, as well as the CEBP-1 transcription factor previously implicated in axon regeneration and cilia maintenance; this induction is mediated by DAF-19. Consistently, both CEBP-1 and the DLK-1 MAPKKK also linked to axon regeneration are necessary for robust cilia regrowth. Finally, we show that cilia truncation and regeneration dynamics vary across neuron types and are modulated by glia. Our results establish that both cell-intrinsic and extrinsic mechanisms promote robust neuronal cilia regeneration in the adult, and that variations in these pathways contribute to neuron type-specific differences in cilia regenerative capacities.

*Chlamydomonas* flagella elongate to half their original length following truncation if new protein synthesis is inhibited indicating that pre-existing ciliary building blocks are sufficient to support partial flagella regrowth in the absence of increased flagellar gene expression ^69^. In contrast, we find that loss of *daf-19* abolishes cilia elongation in *C. elegans,* suggesting that DAF-19-dependent transcriptional changes are essential for all cilia regrowth in this organism. As in *Chlamydomonas* ^73,95,96^, we observed distinct temporal patterns of ciliary gene expression during cilia truncation and regeneration in *C. elegans* such that *osm-6* is upregulated prior to initiation of cilia regrowth. Recent cryo-EM analyses have shown that OSM-6/IFT52 forms a central scaffold of the IFT-B1 complex ^76^. Early upregulation of the OSM-6 core component upon cilia truncation may promote the ordered assembly of pre-existing IFT complexes to support efficient cilia regeneration. In support of this hypothesis, we also observed marked upregulation of OSM-6 but not other examined IFT proteins, in IFT *lof* mutants. In the future, it will be important to identify the entire complement of genes whose expression is altered during cilia truncation and/or regrowth in a neuron type-specific manner, and to determine whether these transcriptional responses are mediated solely by DAF-19 or also by other factors across neuron types.

How might ciliary gene expression be regulated on distinct timescales during cilia disassembly and regeneration? In neurons, activity-dependent and calcium-regulated gene expression occurs in distinct temporal waves driven in part via the early induction of transcription factors which subsequently regulate delayed expression of downstream target genes ^97–99^. Calcium has also been implicated in the regulation of flagellar gene expression and regeneration in *Chlamydomonas* ^100,101^. We speculate that cilia truncation triggers a similar transcriptional cascade with different factors cooperating with, or regulated by, DAF-19 to modulate ciliary gene expression on different timescales. Consistent with this notion, we show that DAF-19 induces expression of the CEBP-1 transcription factor upon cilia truncation. RFX proteins have been shown to act with transcription factors such as CREB and MEF2C to mediate activity- and calcium-regulated neuronal gene expression ^102–104^. DAF-19 functions may be similarly regulated by interactions with signaling-responsive factors potentially including CEBP-1, and/or by post-translational modifications, to dynamically shape ciliary gene expression during cilia truncation and regeneration ^105,106^.

An unexpected finding was that although DAF-19 is essential for the expression of all known ciliary genes ^43,60,62^, this protein is not required for maintenance of IFT gene expression in adult neurons. In *C. elegans*, the FKH-8 forkhead domain transcription factor acts synergistically with DAF-19 to regulate IFT gene expression in a subset of ciliated sensory neurons, although *fkh-8* mutants exhibit relatively minor defects in cilia morphology or IFT gene expression in amphid channel neurons ^107^. Although it is possible that DAF-19 cooperates with members of other transcription factor families to maintain ciliary gene expression in mature amphid sensory neurons ^108,109^, such factors might have been expected to have been identified from extensive forward genetic screens for cilia-defective mutants ^78,110,111^. Alternatively, ciliary gene loci may remain in an open chromatin state following ciliogenesis, and/or ciliary proteins may be highly stable in mature neurons, thereby reducing the requirement for continuous DAF-19-dependent transcription.

Disruption of the microtubule cytoskeleton is a key trigger for axon regeneration. Microtubule depolymerization activates DLK and downstream transcription factors independently of calcium signaling to promote axon regeneration ^84,86,87^, and recent work has shown that mutations in a β-tubulin subunit increase DLK-1 signaling ^112^. Loss of DLK-1 slows but does not abolish cilia regeneration, indicating that this pathway promotes efficient cilia regeneration but is not essential for regrowth. This pathway may regulate cilia regeneration by modulating tubulin posttranslational modifications, inhibiting depolymerizing kinesins, or regulating endocytosis ^87,89,113^. While cilia disruption in chronic IFT mutants increases ciliary DLK-1 levels in the amphid organ ^88,114^, we observed reduced ciliary localization of DLK-1 in ASH/ASI upon acute cilia truncation. This discrepancy may reflect neuron type-specific patterns of DLK-1 localization, or distinct regulatory mechanisms triggered by acute vs chronic cilia truncation. In contrast, expression of *cebp-1* is upregulated in ASH/ASI soma in both IFT mutants as well as upon acute cilia truncation ^85^ (this work). In IFT mutants, *cebp-1* induction is *dlk-1*-dependent in a subset of amphid ciliated neurons but not in ASH/ASI ^85^. Our results indicate that acute cilia truncation upregulates *cebp-1* in ASH/ASI via DAF-19, highlighting the deployment of distinct, neuron-and context-specific pathways for cilia regeneration. We note that although we observe relatively minor effects on cilia length in *dlk-1* or *cebp-1* mutant adults, we cannot exclude the possibility that these molecules also contribute to efficient cilia elongation during embryogenesis.

The cilia of several sensory neurons in *C. elegans* are intimately associated with specific glia ^78,115^. Amphid sheath glia secrete components of the amphid channel matrix, engulf ciliary fragments and extracellular vesicles produced by sensory neurons, and monitor cilia integrity via direct interactions between glia and ciliary proteins such as FIG-1 and DGS-1, respectively ^91,92,94,116^. We find that disruption of glial-channel interactions slows both cilia truncation and regrowth in ASH, whereas glial loss abrogates cilia regrowth in AWB. There appear to be distinct glial microdomains associated with the glia-embedded distal sensory endings of individual neurons; these microdomains help maintain and shape cilia and sensory ending morphologies ^93,117–120^. Recent work has also shown that during aging, channel neurons protect glia-embedded neurons by inducing the unfolded protein response in amphid sheath glia ^121^, suggesting an intricate interaction among neurons and glia in maintaining cilia and thus neuronal functions in the adult.

Cilia regeneration in *C. elegans* exhibits intriguing parallels with axon regeneration mechanisms described in both worms and other organisms. In both processes, neuron-intrinsic pathways including DLK/CEBP1 signaling and induction of pro-regenerative gene expression, as well as extrinsic pathways such as glial cues contribute to structural and functional recovery in a neuron-specific manner ^29,30,33,35,122^. Moreover, axon regeneration is mediated by mechanisms that are partly distinct from those employed during developmental axon outgrowth. Similarly, our results suggest that cilia regrowth may also be driven in part by regeneration-specific programs. Further characterization of the pathways that promote or inhibit cilia regeneration may enable restoration of cilia function in postmitotic cells following injury thereby ameliorating phenotypes associated with acute cilia loss in the adult.

## Acknowledgements

We thank the *Caenorhabditis* Genetics Center and the National BioResource Project (Japan) for strains, the Brandeis Light Microscopy Facility (RRID:SCR_025892) and Andrew Stone for assistance with microscopy, the Brandeis Materials Research Science and Engineering Center (MRSEC) for access to the microfabrication facility, and Ashish Maurya, Priya Dutta, Revanth Sudhireddy and Dom Rinaldi for technical assistance. We are grateful to members of the Sengupta lab, Max Heiman, and Yishi Jin for advice and input. This work was funded in part by the NIH (R35 GM122463 and R21 DC021440 – P.S., T32 GM139798 – S.L.), and a grant from the Rhode Island College Committee for Faculty Scholarship and Development (A.P.). Instrumentation in the Brandeis Light Microscopy Facility was partly funded by a Massachusetts Life Sciences Center Research Infrastructure Program award.

## Author Contributions

Conceptualization, K.J., A.P., P.S.; Methodology, K.J., A.P.. S.N., S.L., Y-M.L.; Investigation, K.J., A.P.. S.N., S.L., L.G., Y-M.L.; Writing – Original Draft, K.J., A.P., P.S.; Writing – Review & Editing, K.J., A.P., S.N., S.L., L.G., Y-M.L., P.S.; Visualization, K.J., A.P., S.L., P.S.; Supervision, P.S., Funding acquisition – P.S.

## METHODS

### C. elegans genetics

*C. elegans* strains were cultured on nematode growth medium (NGM) plates seeded with *Escherichia coli* OP50. The wildtype strain was Bristol, strain N2. All strains were generated using standard genetic techniques. Mutations were verified through PCR-based amplification and/or DNA sequencing. Experimental plasmids were co-injected with either *unc-122*p::GFP or *unc-122*p::dsRed at a concentration of 40 ng/μl to establish transgenic lines. For each injected construct, animals from at least two independent transgenic lines were assessed, with one or two representative lines analyzed in detail. The same extrachromosomal array was examined in both wildtype and mutant backgrounds for comparison. All experiments were conducted using one-day-old adult hermaphrodites unless specified otherwise. All strains used in this work are listed in Table S1>.

### *C. elegans* genome-engineered strains

Endogenously tagged strains were generated using CRISPR/Cas9 genome engineering with Cas9 protein, tracrRNA, and crRNAs synthesized using the EnGen sgRNA Synthesis Kit (NEB #E3322S) (*oy218* and *oy210*) or obtained from IDT, and injected as ribonucleoprotein complexes as previously described ^123^. Repair templates included ssODNs or donor PCR products as indicated in Table S2>. Insertion sequences were added immediately before the stop codon (C-terminal tags) and confirmed by sequencing. Reagents used for genome editing are listed in Table S2>.

### Molecular Biology

The *dlk-1L* cDNA was a kind gift from Yishi Jin. The *sra-6*p*::dlk-1L* (PSAB1424), *sra-6*p*::TIR1::SL2::wrmScarlet* (PSAB1422), and *sra-6*p*::eBFP* (PSAB1425, gift from Nikhila Krishnan) plasmids were generated using standard cloning methods. To clone the *daf-19d* genomic region, sequences from 10166140-10157308 on LG II (Wormbase release WS298) were amplified from N2 genomic DNA and inserted into a derivative of pBluescript (pLink10) via Gibson cloning to generate PSAB1426. All constructs were verified by sequencing.

### Osmotic avoidance assays

Osmotic avoidance assays were performed as previously described ^59^. Briefly, adult worms were first transferred to an NGM plate without food for 5 mins. 10 young adult hermaphrodites were then picked without food into the center of an 8M glycerol ring visualized with xylene cyanol (Sigma-Aldrich X4126). Animals were allowed to move freely on the plate, and the number of animals remaining within the ring were quantified after 10 mins.

### Calcium imaging

Stimulus-evoked imaging of intracellular calcium dynamics in ASH soma was performed using custom microfluidics devices as described previously ^56,124^. Images were acquired using an Olympus BX52WI microscope with a 40x oil objective and Hamamatsu Orca CCD camera at 4 Hz with 4 x 4 binning. Glycerol was diluted in filtered S-basal buffer, and wildtype and mutant worms were paralyzed in 10mM (-)-tetramisole hydrochloride (Sigma L9756) for several mins prior to loading into the microfluidics devices. The 1M glycerol stimulus was prepared fresh prior to each day of imaging, and mutant animals were imaged alongside wildtype controls in each independent experiment. Glycerol-evoked calcium transients in ASH were recorded for 2 cycles of 30 sec S-basal buffer/30 sec 1M glycerol/30 sec S-basal buffer. Data shown are recorded from the second stimulus pulse, as responses to the first pulse are variable. The 0 sec time point for glycerol responses represents the 90 sec following the start of calcium imaging.

Fluorescence changes were analyzed using a custom FIJI script ^56,125^ Images were first aligned using the Template Matching plugin, and manually drawn ROIs were used to identify soma and background regions. The background fluorescence was subtracted from that in the soma, and these values were used for further analysis in RStudio. RStudio was used to calculate the response mean and SEM, and F_0_ was calculated from the average of ΔF/F_0_ value for 5 sec prior to odor onset.

### Auxin experiments

4 mM auxin containing plates were made by dissolving 400mM Auxin (1-naphthaleneacetic acid, Sigma 317918) in 100% ethanol and then adding to NGM. Control plates contained an equal volume of ethanol alone. Plates were seeded with 200 μl *E. coli* OP50, stored at 4 °C and wrapped in aluminum foil to prevent light exposure. For experiments in which cilia lengths were measured, one day-old adult animals were placed on auxin or control ethanol plates for at least 30 mins prior to temperature upshift or downshift. For experiments in which reporter fluorescence intensities were measured, one day-old adult animals were placed on auxin or control ethanol plates overnight prior to temperature upshift or downshift.

### Dye-filling

Stock solutions of 1,1′-dioctadecyl-3,3,3′,3′-tetramethylindocarbocyanine perchlorate (DiI) at 1 mg/ml in *N,N*-dimethylformamide (Sigma-Aldrich 468495) were stored at -20°C. To evaluate dye uptake, animals were immersed in 0.1mg/ml dye for one hour before examination.

### Fluorescent reporter imaging and image analysis

One day-old adult hermaphrodites were immobilized using 10mM tetramisole hydrochloride (Sigma-Aldrich L9756) and mounted on 10% agarose pads prepared on microscope slides with coverslips. Animals were imaged on an Andor CSU-W1 spinning disc confocal microscope with a Leica DM6000b body using a 100X objective. Images were acquired using Andor Fusion software version 2.4.0.14. Additional imaging was performed on a 3i Marianas spinning disc confocal microscope with a Zeiss Observer Z1 body equipped with a Yokagawa CSU-X1 spinning disk confocal head using a 100x objective. Images were acquired using Slidebook software. *z*-stacks were acquired with a step-size of 0.2 µm. For Figures 3D, 3E and 5C animals were imaged on an Andor Dragonfly 620SR spinning disc confocal microscope with a Nikon Ti2 body with an ASI Piezo Z-stage using a 60x objective. Images were acquired using Andor Fusion software version 2.7.0. *z*-stacks were acquired with a step-size of 0.2 µm.

Imaging parameters were kept consistent between wildtype and mutant samples. For cilia length measurements, image processing and analysis was conducted using FIJI/ImageJ (NIH). ASH cilia were identified as expressing *sra-6*p*::gfp* at higher levels than in neighboring ASI cilia. For fluorescent reporter intensity measurements, cell bodies or cilia were rendered in 3D by absolute intensity thresholding using the Surface tool in Imaris 10.2 (Oxford Instruments), prior to plotting sum intensity values. Since ASH and ASI neurons could not be differentiated by reporter expression levels, measurements included cilia or soma of either or both neurons.

### IFT Measurements

Intraflagellar transport (IFT) analyses were carried out as described previously ^56^. In brief, time-lapse recordings of fluorescently tagged IFT proteins in ASH cilia were carried out on a 3i Marianas spinning disc confocal microscope with a Zeiss Observer Z1 body equipped with a Yokagawa CSU-X1 spinning disk confocal head (Figure 1G) or an Andor CSU-W1 spinning disc confocal microscope with a Leica DM6000b body (Figure S4C). Images were captured over a 30 sec time frame at 250 msec exposure intervals and at 100X magnification. In FIJI/ImageJ, a line was manually drawn from the base through the middle segment of ASH cilia. Kymographs were generated using the Multi Kymograph and Template Matching plugins (ImageJ) to account for motion-related artifacts. After manually tracking IFT particles, velocity measurements were obtained by tracing line segments along each individual IFT trajectory.

### Statistical analyses

All graphs were generated and analyzed using RStudio (for calcium imaging analysis) and Prism 9 software. Comparisons among strains were performed using an unpaired Welch’s t-test or one-way ANOVA followed by Tukey’s multiple comparisons test. The number of animals examined in each experiment, significance values, and number of independent experiments are provided in the figure legends.

---

**Figure S1.**
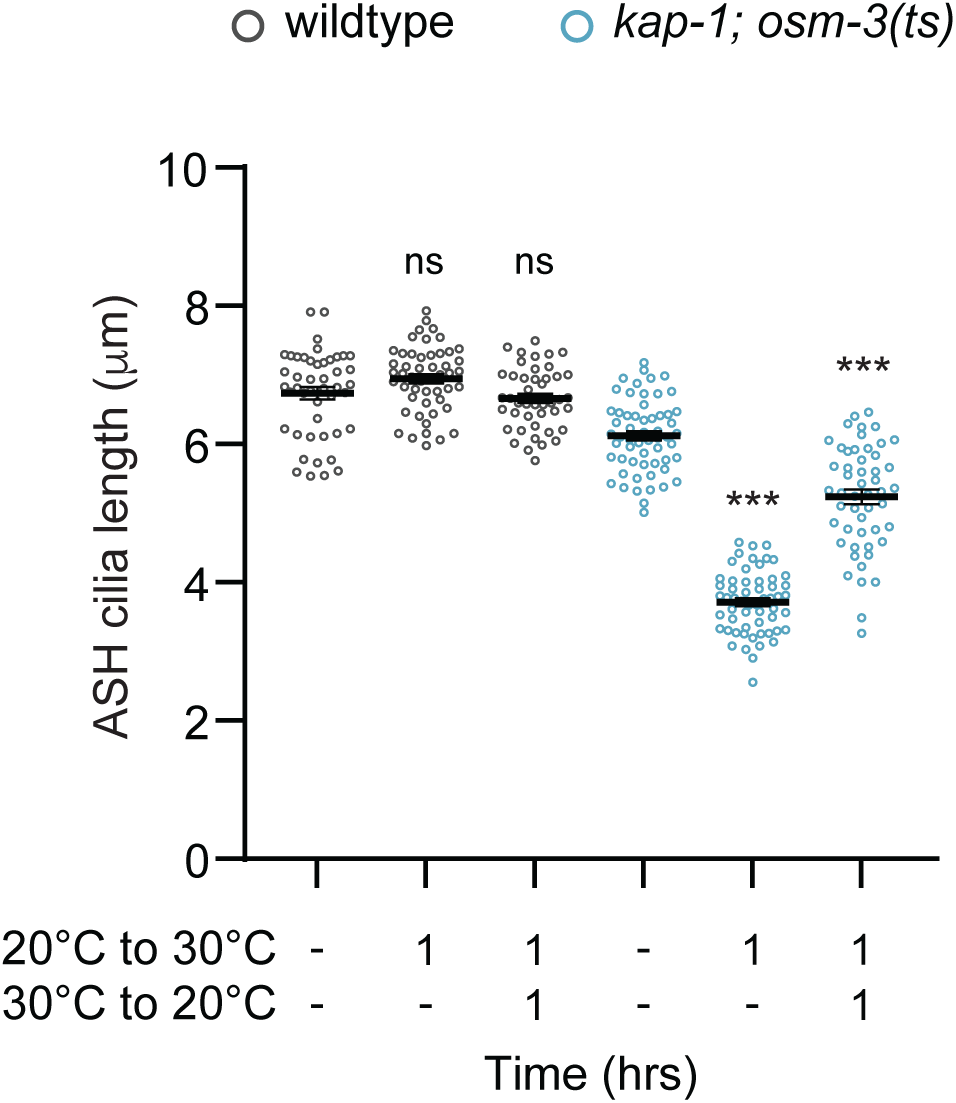
related to Figure 1. ASH cilia truncate and regrow rapidly in the adult. Quantification of ASH cilia length in wildtype (black) and *kap-1(ok676); osm-3(oy156ts)* (blue) mutants at the indicated temperature shift conditions. Each circle is the length of a single ASH cilium. ***: different at *P*<0.001 from unshifted within each genotype; ns: not significant (one-way ANOVA with Tukey’s multiple comparison test). Errors are SEM. n≥44 each; 3 independent experiments.

**Figure S2.**
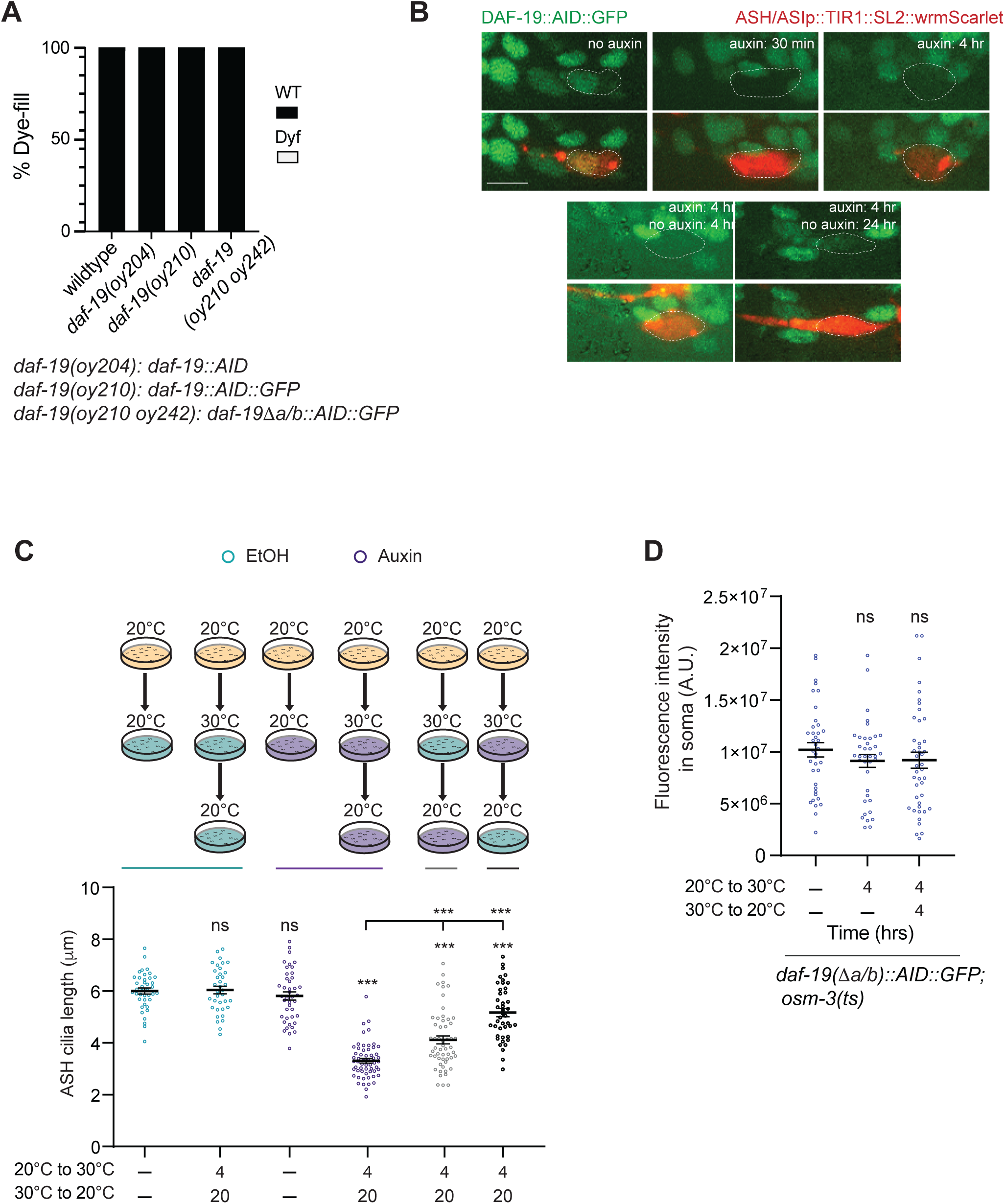
related to Figures 2 and 3. DAF-19 is required for cilia regeneration but is dispensable for maintenance. **A)** Percentage of animals of the indicated genotypes in which a subset of ciliated neurons including ASH/ASI are filled with dye. n>300 animals each; 3 independent experiments. Dyf: dye-filling defective. **B)** Representative images of endogenously tagged DAF-19::AID::GFP expression in ASH/ASI neurons in the absence or presence of auxin as indicated. These neurons also express *sra-6*p*::TIR1::SL2::wrmScarlet* from an extrachromosomal array. ASH/ASI soma are indicated with dashed outlines. Scale bar: 5 μm. **C)** (Top) Schematic of experimental conditions for ASH cilia length quantifications (bottom). Each circle is the length of a single ASH cilium. ***: different at *P*<0.001 from unshifted or between indicated; ns: not significant (one-way ANOVA with Tukey’s multiple comparison test). Errors are SEM. n≥36 each; 3 independent experiments. **D)** Quantification of reporter fluorescence intensity in ASH/ASI soma of endogenously reporter-tagged *daf-19(oy210 oy242Δa/b)* in *osm-3(oy156ts)* animals in the shown temperature shift conditions. Each circle is the value from an ASH and/or ASI soma in one amphid organ. ns: not significant from unshifted (one-way ANOVA with Tukey multiple comparison’s test). Errors are SEM. n≥38 each; 2 independent experiments.

**Figure S3.**
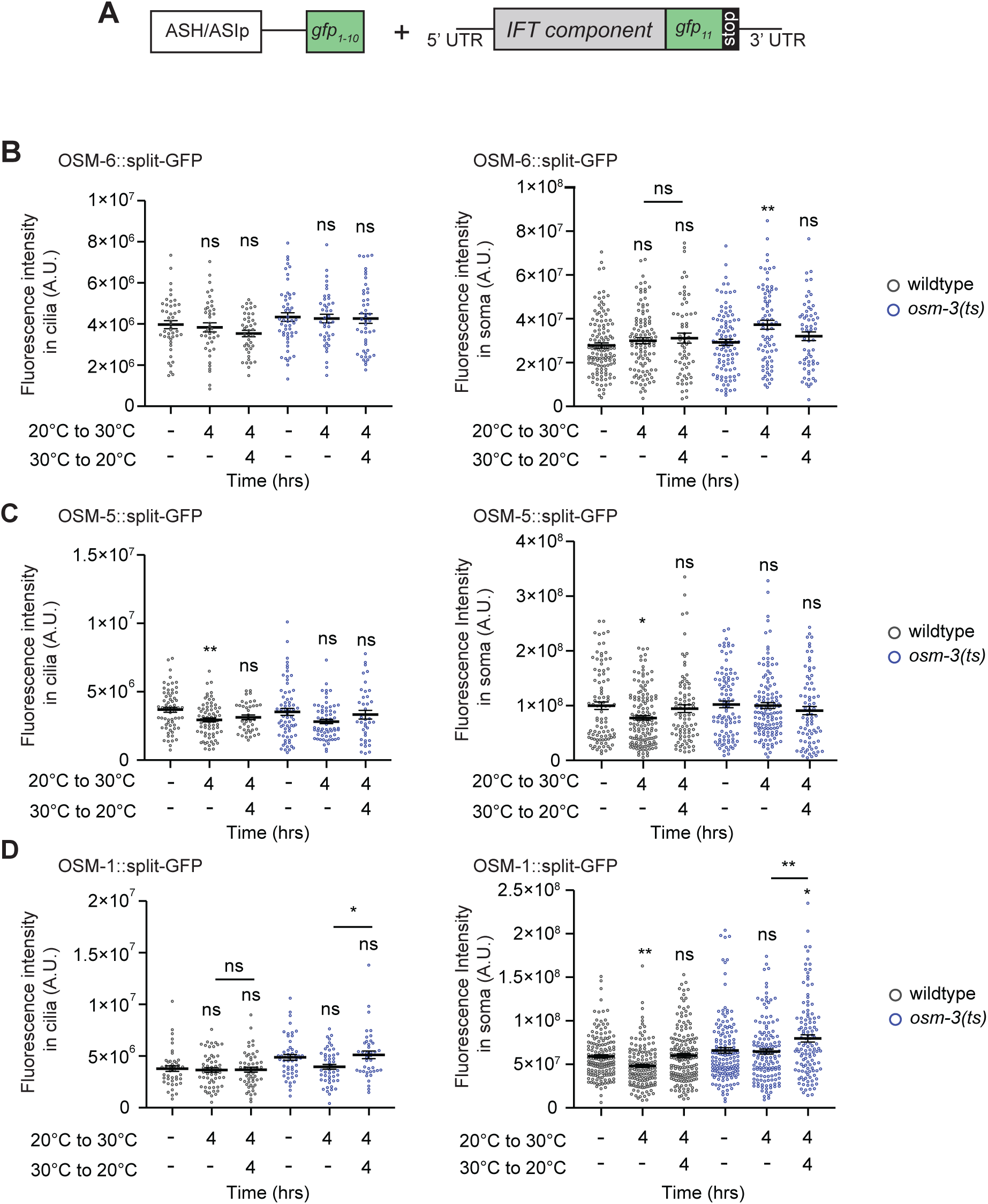
related to Figure 4. IFT protein levels in ASH soma are altered upon cilia truncation and regeneration. **A)** Schematic of endogenously tagged translational reporter expression. **B-D)** Quantification of fluorescence intensities in ASH/ASI cilia (left) and soma (right) of wildtype (black) or *osm-3(oy156ts)* (blue) animals expressing the indicated IFT translational reporters in the shown temperature shift conditions. Each circle is the value from an ASH and/or ASI cilia (left) or soma (right) in an amphid organ. * and **: different at *P*<0.05 and 0.01, respectively, from unshifted within each genotype or between indicated; ns: not significant (one-way ANOVA with Tukey’s multiple comparison test or unpaired Welch’s t-test). Errors are SEM. n≥39 each; >3 independent experiments.

**Figure S4.**
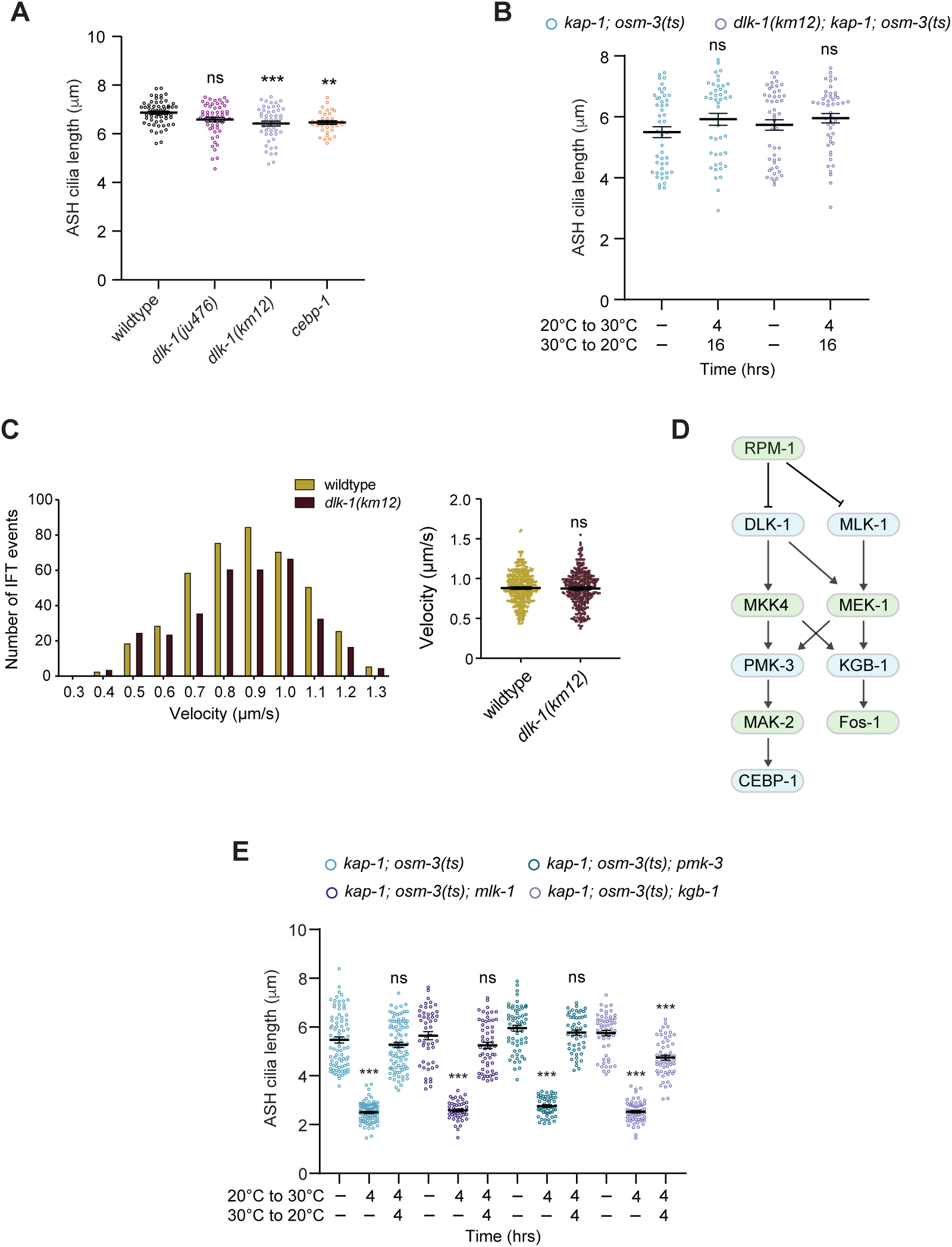
related to Figure 5. DLK-1 and CEBP-1 are necessary for efficient cilia regeneration. **A)** Quantification of ASH cilia length in the shown genetic backgrounds Each circle is the length of a single ASH cilium. ** and ***: different at *P*<0.01 and 0.001 from wildtype; ns: not significant (one-way ANOVA with Tukey’s multiple comparison test). Errors are SEM. n≥35 each; 2 independent experiments. **B,E)** Quantification of ASH cilia length in the shown genetic backgrounds at the indicated temperature shift conditions. Each circle is the length of a single ASH cilium. ***: different at *P*<0.001 from unshifted within each genotype; ns: not significant (B: unpaired Welch’s t-test, E: one-way ANOVA with Tukey’s multiple comparison test). Errors are SEM. n≥44 each; 3 independent experiments. **C)** Histogram (left) and quantification (right) of velocities of reconstituted endogenously tagged OSM-6::split-GFP in ASH cilia in wildtype and *dlk-1(km12)* animals. n≥24 animals per genotype; 3 independent experiments. ns: not significant (unpaired Welch’s t-test). **D)** Known DLK-1-regulated signaling pathways. Adapted from ^1,2^.

**Figure S5.**
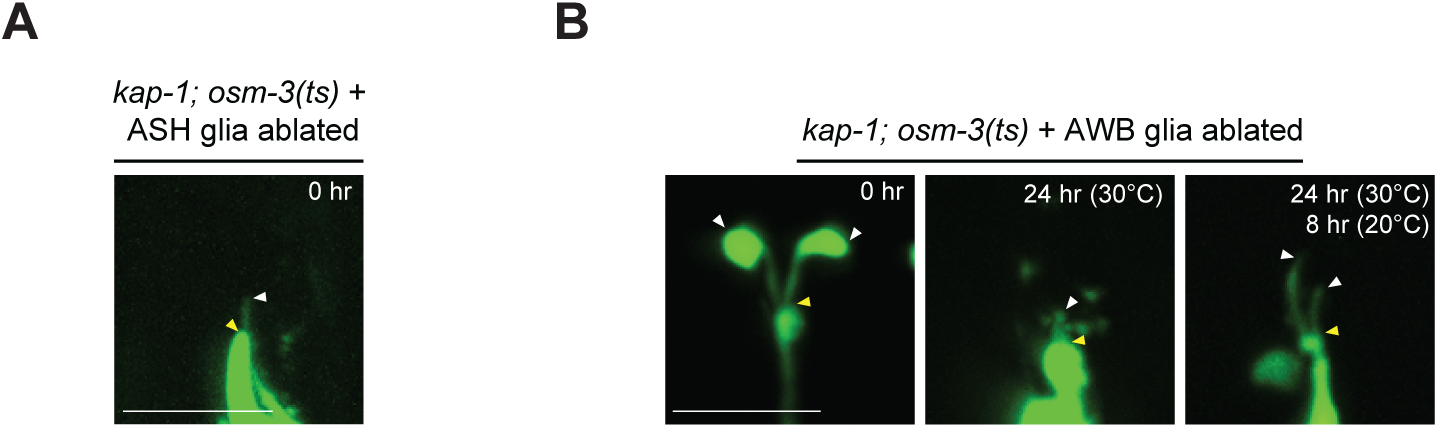
related to Figure 6. Cilia morphology is altered in the absence of glia. Representative images of ASH (A) and AWB (B) cilia in the indicated genetic backgrounds and temperature shift conditions. Yellow/white arrowheads: cilia base/cilia tip. Scale bars: 5 μm.

**Table S1.**
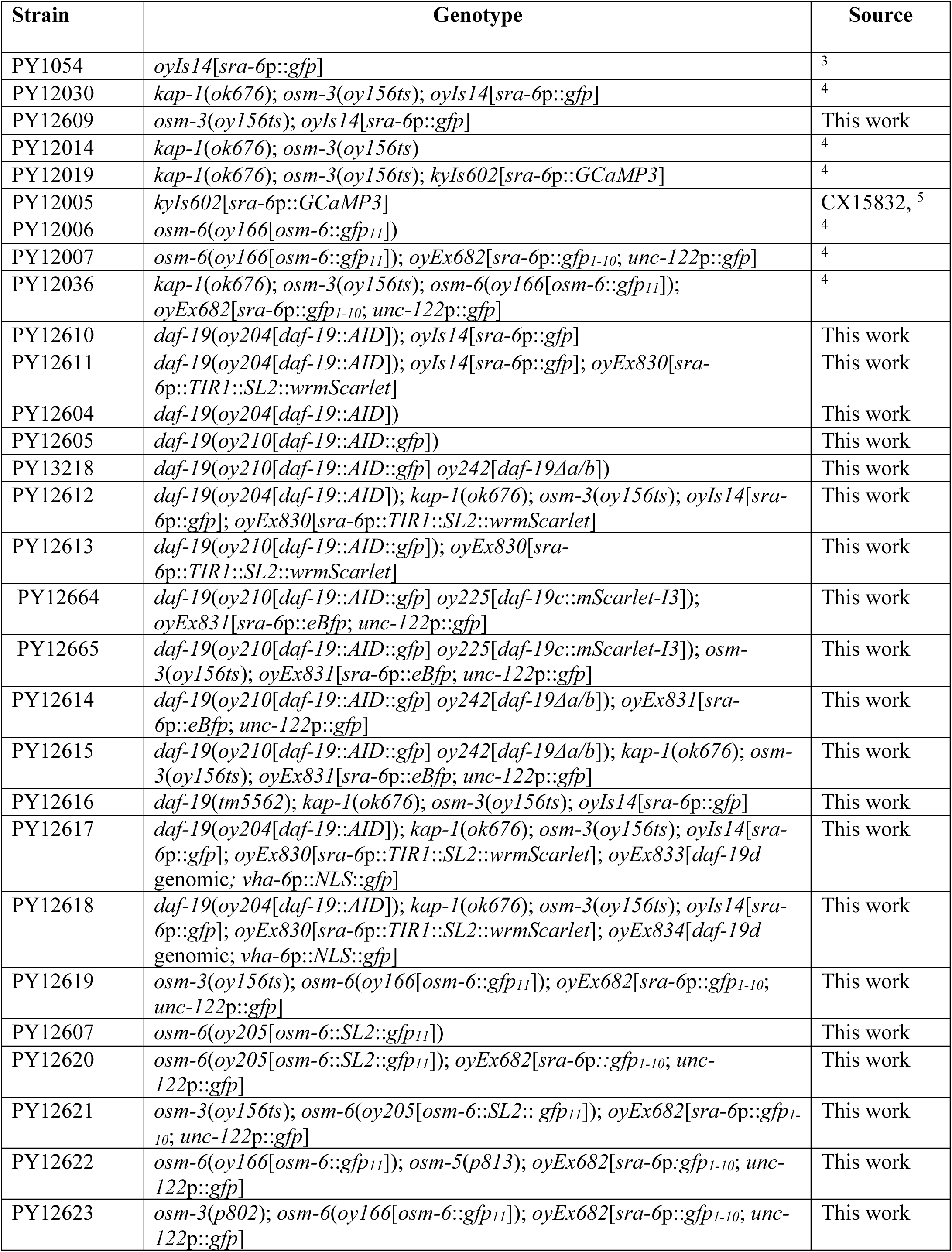
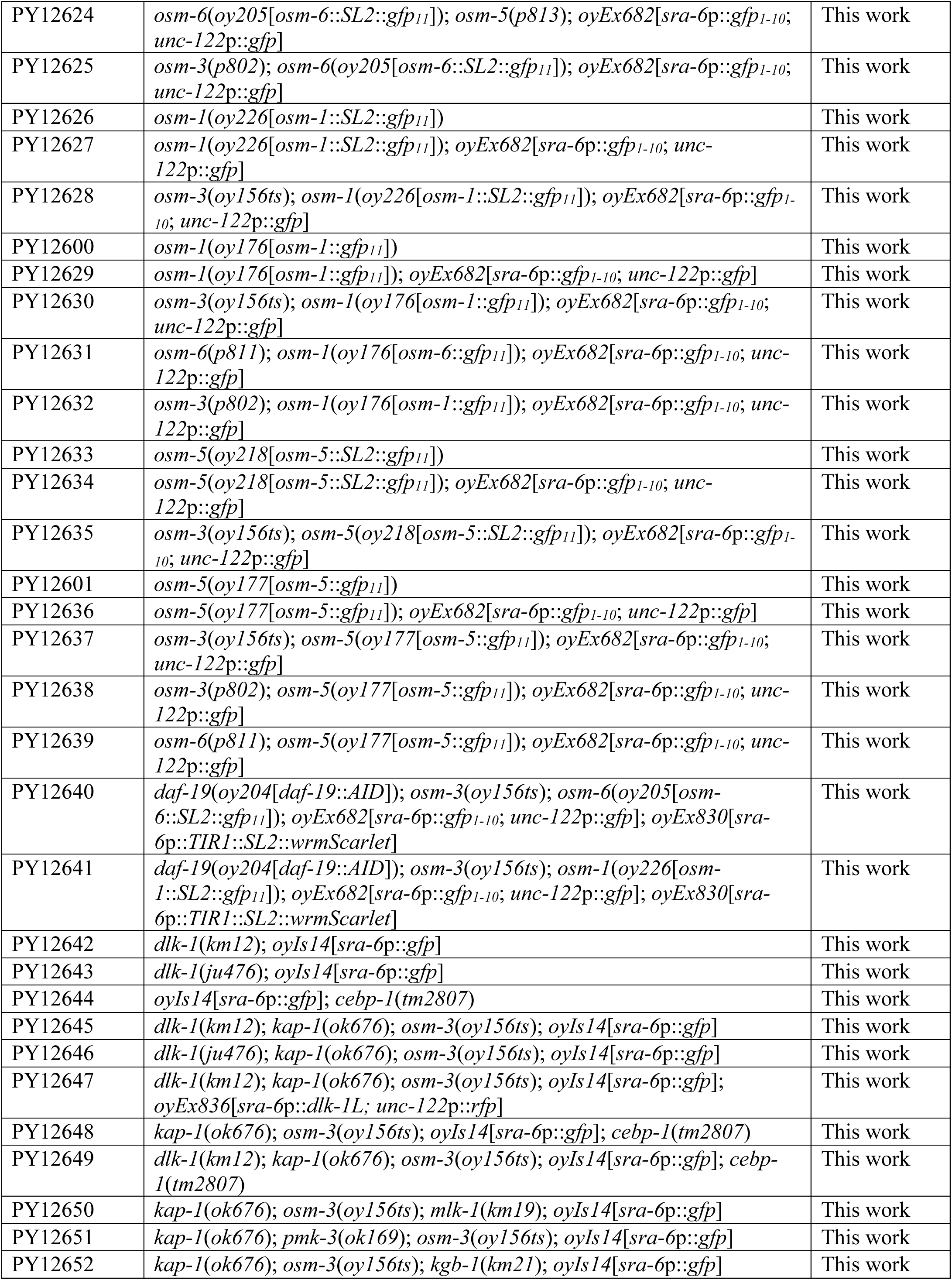
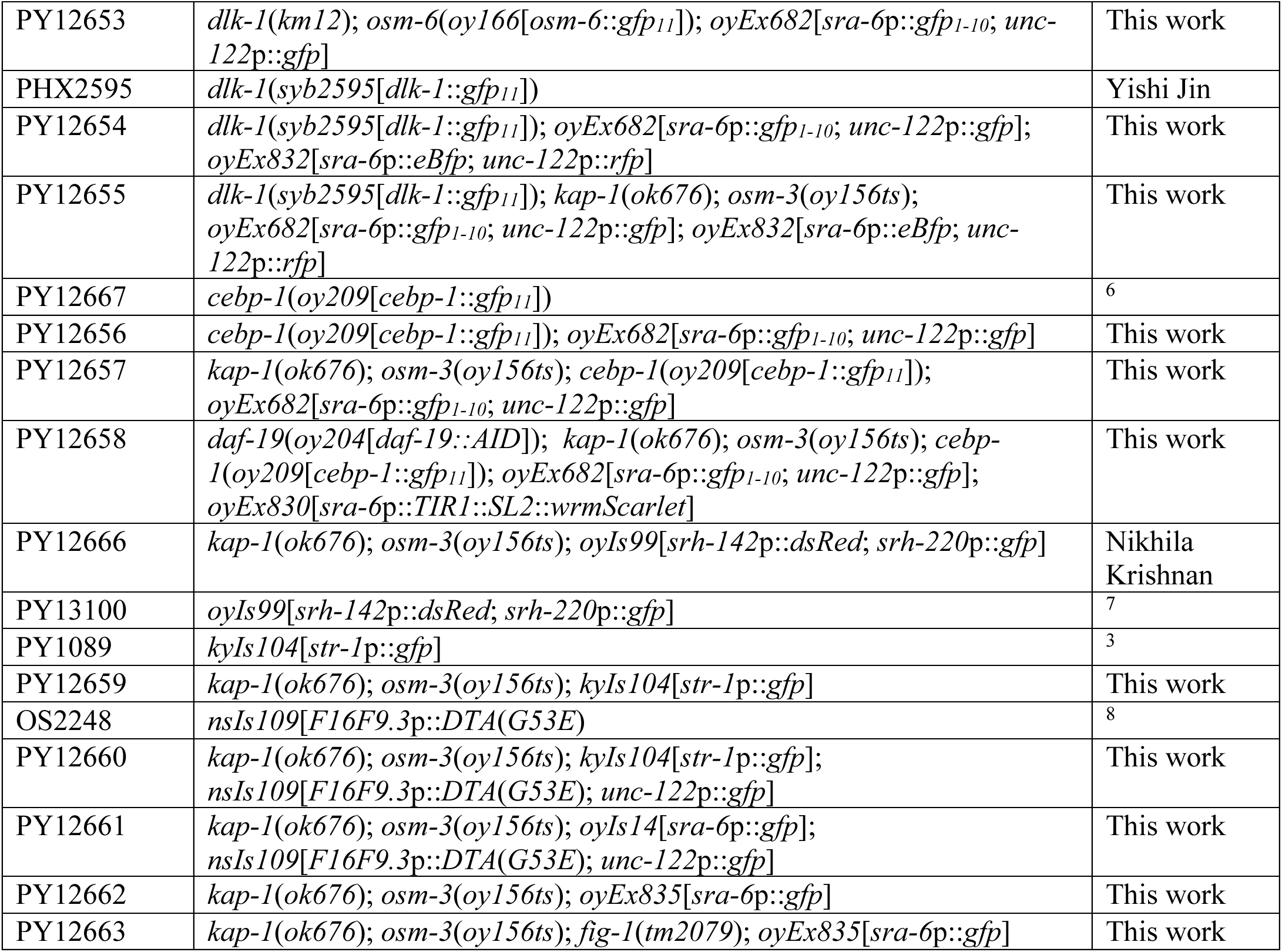
related to all Figures. List of strains used in this work.

**Table S2.**
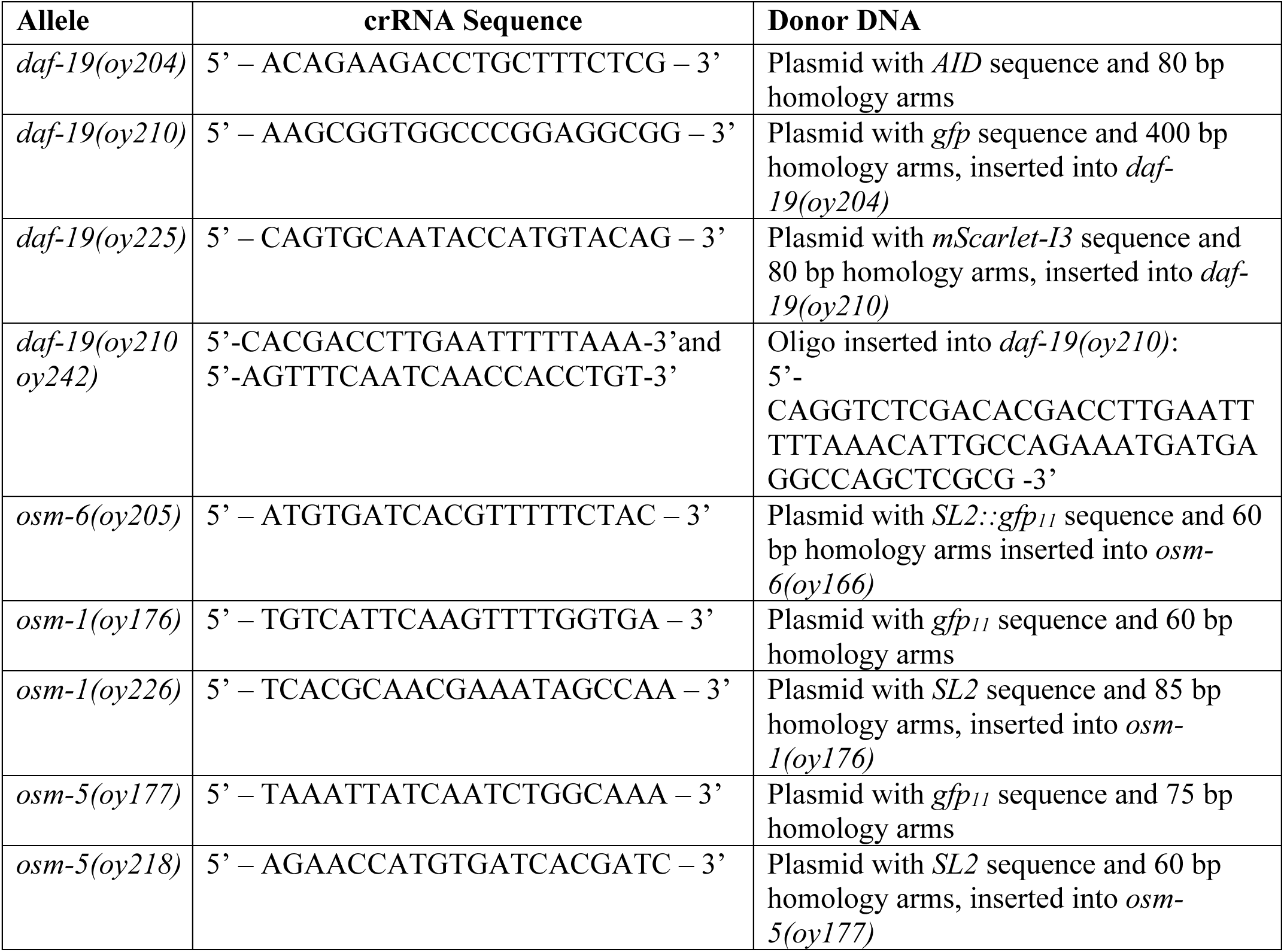
related to all Figures. Gene editing reagents used in this work.

